# Somatic uniparental disomy mitigates the most damaging *EFL1* allele combination in Shwachman-Diamond syndrome

**DOI:** 10.1101/483362

**Authors:** Sangmoon Lee, Chang Hoon Shin, Jawon Lee, Seong Dong Jeong, Che Ry Hong, Jun-Dae Kim, Ah-Ra Kim, Soo Jin Son, Oleksandr Kokhan, Taekyeong Yoo, Jae Sung Ko, Young Bae Sohn, Ok-Hwa Kim, Jung Min Ko, Tae-Joon Cho, Nathan T. Wright, Je Kyung Seong, Suk-Won Jin, Hyoung Jin Kang, Hyeon Ho Kim, Murim Choi

**Author notes:** H.J. Kang, H.H.Kim, and M.Choi contributed equally to this paper Correspondence to Murim Choi; or Hyeon Ho Kim or Hyoung Jin Kang. S. Lee, and C.H. Shin contributed equally to this paper. Sangmoon Lee (Department of Neurosciences, University of California, San Diego, La Jolla, CA 92093, USA). Chang Hoon Shin (Laboratory of Genetics and Genomics, National Institute on Aging Intramural Research Program, National Institutes of Health, Baltimore, MD 21224, USA).

## Abstract

Shwachman-Diamond syndrome (SDS; OMIM: #260400) is caused by variants in *SBDS* (Shwachman-Bodian-Diamond syndrome gene), which encodes a protein that plays an important role in ribosome assembly. Recent reports suggest that recessive variants in *EFL1* are also responsible for SDS. However, the precise genetic mechanism that leads to *EFL1*-induced SDS remains incompletely understood. Here we present three unrelated Korean SDS patients that carry biallelic pathogenic variants in *EFL1* with biased allele frequencies, resulting from a bone marrow-specific somatic uniparental disomy (UPD) in chromosome 15. The recombination events generated cells that were homozygous for the relatively milder variant, allowing for the evasion of catastrophic physiological consequences. Still, the milder *EFL1* variant was solely able to impair 80S ribosome assembly and induce SDS features in cell line, zebrafish, and mouse models. The loss of *EFL1* resulted in a pronounced inhibition of terminal oligo-pyrimidine element-containing ribosomal protein transcript 80S assembly. Therefore, we propose a more accurate pathogenesis mechanism of EFL1 dysfunction that eventually leads to aberrant translational control and ribosomopathy.

## Introduction

Patients clinically diagnosed with Shwachman-Diamond syndrome (SDS; OMIM: #260400) present with a constellation of disorders, such as hematologic manifestations, exocrine pancreatic dysfunction with fatty infiltration, and skeletal dysplasia that results in short stature^1–3^. The hematologic manifestations include neutropenia or, less severely, thrombocytopenia and anemia with a predisposition for myelodysplastic syndrome and acute myeloid leukemia transformations^4,5^. Variants in *SBDS* (Shwachman-Bodian-Diamond syndrome gene), which encodes a protein that plays an important role in ribosome assembly, are mainly responsible for the disease^1,6–8^. Thus, SDS is considered to be a ribosomopathy, which is a collective term that is used to describe a group of congenital disorders caused by problems in ribosome biogenesis, assembly or function^9^. Moreover, ~10% of clinically diagnosed SDS cases do not contain any pathogenic *SBDS* variants, suggesting the existence of additional genetic mechanisms that lead to the disorder^1,6^. Recent reports demonstrate that variants in genes other than *SBDS*, namely *EFL1*, *DNAJC21,* and *SRP54*, are implicated with bone marrow failure syndrome and SDS^10–15^. Homozygous variants of *EFL1* cause an SDS-like syndrome in a recessive manner, which is highlighted by the observation that three of the seven reported kindreds underwent consanguineous marriages^10,11,15^. EFL1 directly interacts with SBDS to release eukaryotic translation initiation factor 6 (eIF6) from the 60S ribosomal subunit for 80S ribosomal assembly^7,16^. Thus, it needs to be further investigated whether there is an additional genetic mechanism that leads to SDS in outbred populations other than through homozygous pathogenic variants in *EFL1*.

As more human genomes with or without clinical significances continue to be sequenced, it has become clear that variants of unknown significances (VUS) pose a substantial obstacle in the interpretation of genotype-phenotype relationships. As many variants are believed to possess the ability to cause alternations at the molecular level but only with sub-clinical levels of severity, numerous scenarios that enable VUS to acquire clinical significances have been postulated. One of these scenarios involves the assessment of somatically acquired uniparental disomy (UPD) in the hematopoietic system; although only a small number of instances have been previously reported. Notable examples include myeloid neoplasia^17,18^, immunodeficiency^19^, and a single case of sickle cell disease^20^.

In this study, we demonstrated a disease-causing mechanism in patients who inherited compound heterozygous variants in *EFL1*. A mosaic UPD caused a loss-of-heterozygosity (LOH) in the *EFL1* locus in the bone marrow and blood, simultaneously homozygosing the less damaging variant and decreasing the representation of the more damaging variant to avoid worse hematologic phenotypes. However, this still led to EFL1 dysfunction in the bone marrow and resulted in SDS features. We further demonstrated that the remaining variant by the UPD was a hypomorph and pathogenic, by investigating the molecular mechanism of the EFL1 dysfunction in cell and animal models. Therefore, searching for a pathogenic variant that was caused by a non-conventional pathway may increase the probability of identifying the genetic cause and improve our understanding of the disease mechanism, and this approach could possibly benefit additional patients with severe hematological abnormalities.

## Results

### Shwachman-Diamond syndrome patients without *SBDS* variants

We recruited three unrelated and non-consanguineous Korean SDS patients without plausible recessive mutations in *SBDS* (Fig. 1a; Table S1; Supplemental Clinical Narratives). Proband I-1 is a 3-year-old boy who had severe intrauterine growth retardation that resulted in a preterm delivery (35+3 weeks) and a birth weight of 1.7 kg. He had thrombocytopenia, neutropenia, and anemia at 2-months of age. A bone marrow examination performed at 6-months of age revealed hypocellularity, reduced megakaryocytes, and an increase in iron storage. He also had pancreatic lipomatosis, along with an exocrine pancreatic insufficiency, and metaphyseal chondrodysplasia (Fig. 1a; Fig. S1). Proband II-1 is a 9-year-old female who had severe intrauterine growth retardation that resulted in a preterm delivery (36 weeks) and a birth weight of 1.6 kg. She had a diffuse fatty infiltration of the pancreas and metaphyseal chondrodysplasia that was accompanied by osteopenia and short stature (Fig. 1a). Her sister was unaffected. Proband III-1 is a 25-year-old male who did not have any perinatal problems except for a low birth weight (40+4 weeks, 2.4 kg). At 2-years-old, he had pancreatic exocrine and endocrine insufficiencies, thrombocytopenia, anemia, intermittent neutropenia, metaphyseal chondrodysplasia, and ichthyosis. Later, he developed osteoporosis, hepatomegaly, and a total fatty change of the pancreas (Fig. 1a).

**Fig. 1.**
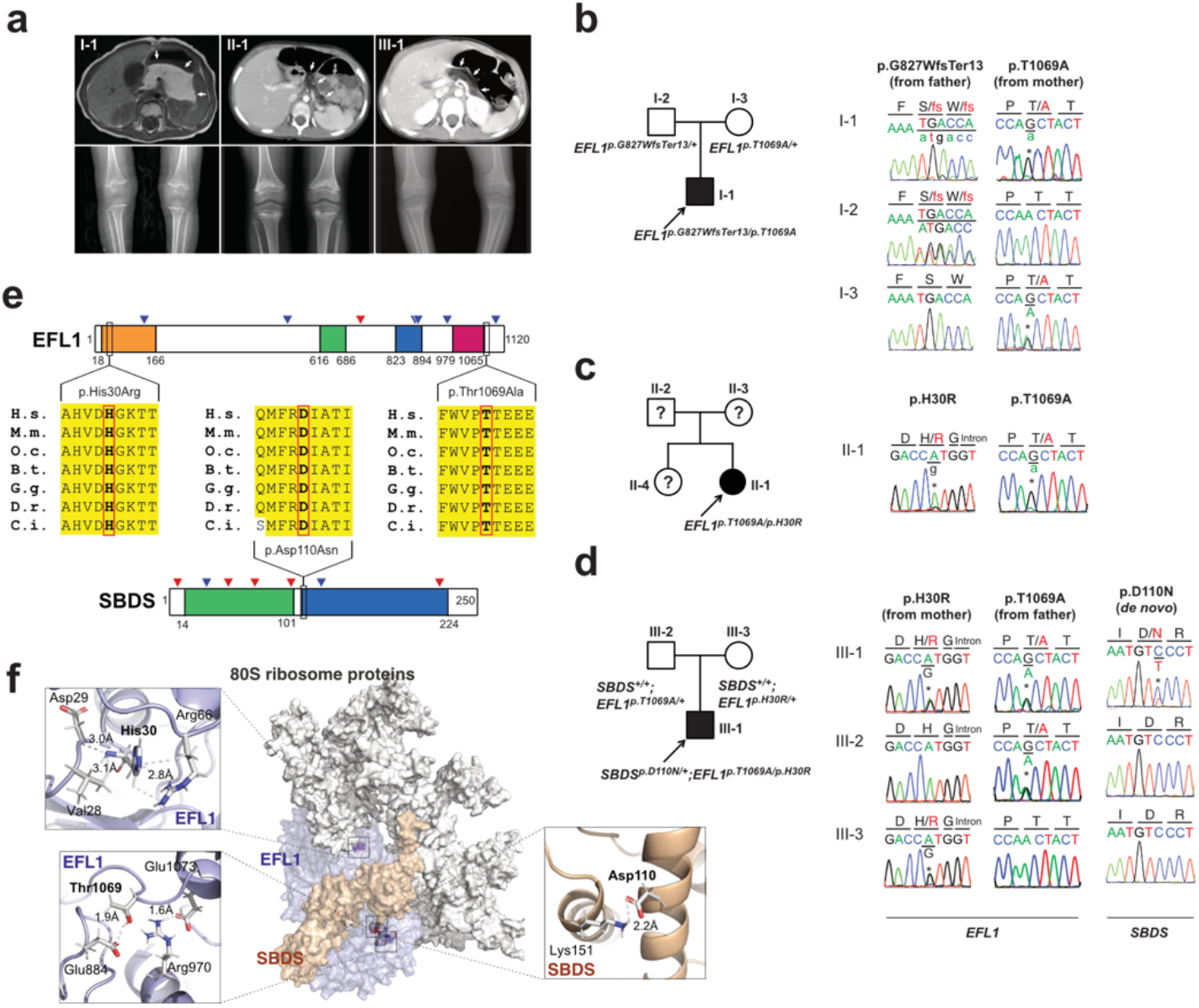
*EFL1* variants in *SBDS*-negative SDS patients. **a** Non-contrast T1-weighted abdominal MRI (I-1) or CT images (II-1 and III-1) showing a diffuse enlargement with lipomatosis of the pancreas (arrows) (upper) and both knees with metaphyseal widening and irregularities in the femora in the three patients and associated genu varum in III-1 (lower). **b-d** Pedigrees and Sanger sequencing traces showing the inherited *EFL1* variants and the *de novo SBDS*^*p.Asp110Asn*^ variant in the three families. DNA was extracted from whole blood samples. Note that the lower case nucleotide letters were used on top of the Sanger traces to reflect their minor representation in the patient samples. **e** The residues with nonsynonymous changes in EFL1 and SBDS are evolutionarily conserved. The arrowheads on the protein maps denote previously reported pathogenic variants (Blue: missense, red: LoF). **f** Molecular modeling-based structural analysis using PDB 5ANC^7^. The insets show the detailed interactions involving EFL1 His30, Thr1069 and SBDS Asp110. B.t., *Bos taurus*; C.i., *Ciona intestinalis*; D.m., *Drosophila melanogaster*; D.r., *Danio rerio*; G.g., *Gallus gallus*; H.s., *Homo sapiens*; M.m., *Mus musculus*; O.c., *Oryctolagus cuniculus*.

### Identification of mosaic *EFL1* variants

To identify the genetic factors that predisposed the three patients to SDS, we exome-sequenced the patients and the available parental DNA was extracted from whole blood (Table S2). Notably, the heterozygous p.Thr1069Ala variant of *EFL1* (chr15:82,422,872 T>C, hg19, NM_024580.5:c.3205A>G) was identified in all three patients (Fig. 1b-d, Table 1; Table S3), which was not previously found in SDS patients. Based on gnomAD, it is a low frequency variant that was carried by three individuals among 17,972 alleles in the East Asian population (East Asian allele frequency (EA-AF) = 1.7 x 10^−4^)^21^. Under the assumption that *EFL1* variants function in a recessive manner, we sought to ascertain additional variants that may pose increased damage to the gene function. Remarkably, we found a second variants in *EFL1* in all three patients which were not detected by the initial analysis either due to a low number of variant supporting reads in the proband or due to a low mapping quality caused by the sequence similarity between the *EFL1* and *EFL1P1* loci. Proband I-1 carried a paternally-originated frameshift variant p.Gly827TrpfsTer13 (chr15:82,444,316 C>CA, hg19, NM_024580.5:c.2478dupT, gnomAD EA-AF = 0) with a minor allele frequency (MAF) of 8.3%, and Proband II-1 and III-1 carried an inherited missense variant p.His30Arg (chr15: 82,554,031 T>C, hg19, NM_024580.5:c.89A>G, gnomAD EA-AF = 5.1 x 10^−5^) with MAFs of 14.8% and 36.8%, respectively (Table 1). In addition, Proband III-1 harbored a heterozygous *SBDS* p.Asn110Asp variant as *de novo* (Fig. 1d; Fig. S2), which is never seen in the control databases. Then we noted that the non-reference allele of *EFL1* p.Thr1069Ala for I-1 and II-1 was dominantly covered compared to the reference allele (Table 1), which caused it to function like an incomplete homozygous variant, while the ratio was comparable in III-1 (Fig. 1c; Table 1). All the *EFL1* variants were equally represented in the parental carriers. The patients did not carry any variant in other SDS-associated genes, including *DNAJC21* and *SRP54* (data not shown). The two EFL1 amino acid residues harboring the missense variants (His30 and Thr1069) are highly conserved throughout evolution and are predicted to be pathogenic (Fig. 1e; Table S4). The protein structure analysis suggested that the His30 and Thr1069 residues form hydrogen bonds with neighboring residues, which would presumably confer stability to the protein structure. Notably, previously reported pathogenic residues Cys883 and Arg970 lie close to Thr1069 (Fig. 1f; Fig. S3)^15^. The SBDS Asn110 reside is also conserved among the vertebrate species (Fig. 1e; Table S4) and is expected to interact with neighboring amino acids, including Lys151 (Fig. 1f).

**Table 1.** Coverage depths and allele frequencies of *EFL1* variants

| Patient | Variant in *EFL1* (M: maternal origin, P: paternal, or U: unknown) | Method | Allele coverage depths (ref. allele:non-ref. allele) (% non-ref. allele) |  |  | AF in gnomAD |  |
| --- | --- | --- | --- | --- | --- | --- | --- |
|  |  |  | Patient | Mother | Father | Overall | East Asian |
| I-1 | p.Thr1069Ala (M) | WES | 10:64 (86.5%) | 39:40 (50.1%) | 33:0 (0%) | 3.2 x 10<sup>-5</sup> | 1.7 x 10<sup>-4</sup> |
|  |  | Amplicon seq. | 23,914:131,714 (84.6%) | 63,582:63,954 (50.1%) | 121,502:507 (0.4%) |  |  |
|  | p.Gly827TrpfsTer13 (P) | WES | 22:2 (8.3%) | 27:0 (0%) | 9:14 (60.9%) | 0 | 0 |
|  |  | Amplicon seq. | 243,610:43,042 (15.0%) | 505,158:74 (0.01%) | 221,532:218,976 (49.7%) |  |  |
| II-1 | p.Thr1069Ala (U) | WES | 15:59 (79.7%) | N/A | N/A | 3.2 x 10<sup>-5</sup> | 1.7 x 10<sup>-4</sup> |
|  | p.His30Arg (U) | WES | 127:22 (14.8%) | N/A | N/A | 1.8 x 10<sup>-5</sup> | 5.1 x 10<sup>-5</sup> |
| III-1 | p.Thr1069Ala (P) | WES | 43:30 (41.1%) | N/A | N/A | 3.2 x 10<sup>-5</sup> | 1.7 x 10<sup>-4</sup> |
|  |  | Amplicon seq. | 135,196:149,160 (52.5%) | 189,517:1,786 (0.9%) | 87,347:85,317 (49.4%) |  |  |
|  | p.His30Arg (M) | WES | 74:43 (36.8%) | N/A | N/A | 1.8 x 10<sup>-5</sup> | 5.1 x 10<sup>-5</sup> |
|  |  | Amplicon seq. | 25,316:22,680 (47.3%) | 119,382:121,508 (50.4%) | 41,684:76 (0.2%) |  |  |

To understand the genetic cause of this observation, we investigated whether large-scale structural variants exist that encompass the region. Indeed, all the patients carried a partial LOH in chromosome 15, where *EFL1* resides (Fig. 2a; Figs. S4 and S5). This LOH was copy-neutral and was not seen in the healthy parents (Fig. S6), suggesting that it was caused by a somatic UPD. This LOH of the *EFL1* locus is not frequently found in healthy Korean individuals (1/3,667 = 2.7 x 10^−4^, Fig. S7). Also, according to a survey of hematopoietic chromosomal mosaicism events, 117 of 151,202 apparently normal individuals carry a copy-neutral LOH or a copy-number deletion of the *EFL1* locus (7.7 x 10^−4^)^22^. Thus, the LOH of the *EFL1* locus is a rare event. The sizes of the LOH intervals were variable among the patients (100%, 100% and 27.8% of the entire chromosome span for I-1, II-1, and III-1, respectively; Fig. 2a). We also sought to identify the haplotype origins of the variants that the probands carried and observed that the chromosomes that were dominantly represented (*i.e.*, the maternal chromosome for I-1 and the paternal chromosome for III-1) harbored *EFL1*^p.Thr1069Ala^, which was consistent with their higher coverage ratios compared to other variants as documented in the WES analysis (Fig. 2a). To test if the UPD event occurred in a mosaic pattern and whether LOH-carrying and non-LOH-carrying cells co-exist, a single-cell SNP microarray experiment was performed using bone marrow (I-1) or cells from buccal swabs (III-1). As expected, complete LOHs in chromosome 15 were observed in a subset of the cells, confirming the mosaic UPD events that preferentially selected *EFL1*^p.Thr1069Ala^ over the other variants (*i.e.*, *EFL1*^p.Gly827TrpfsTer13^ and *EFL1*^p.His30Arg^; Fig. 2b-d; Fig. S8). Therefore, cells containing UPD of chromosome 15 are expected to be homozygous for the *EFL1*^p.Thr1069Ala^ allele and be a homozygous reference for the other two alleles. II-1 was not tested, because the sample was unavailable. Next, we checked the spatial extent of the mosaic UPD by subjecting all available tissue samples from I-1 to a high-depth amplicon sequencing analysis (>10,000X coverage depths). Variant AFs of p.Thr1069Ala in I-1 were ~0.85 in the peripheral blood and bone marrow but ~0.5 in most tissues. Conversely, AFs of p.Gly827TrpfsTer13 in the peripheral blood and bone marrow were ~0.15 and displayed complementary frequencies to those of p.Thr1069Ala (Fig. 2e). This observation suggested that the mosaic UPD was restricted at least to the bone marrow. These results were concordant with the Sanger sequencing results (Figs. S9-10). The degree of mosaicism in the bone marrow tissue changed dynamically over the time course of I-1, but did not strongly correlate with the clinical status of the patient (Fig. S11). These results suggest that compound heterozygous variants that may disable *EFL1* function and impair cell survival formed a cellular environment such that cells with (less damaging) recombinant alleles gained survival advantages over the parental ones (Fig. 2f).

**Fig. 2.**
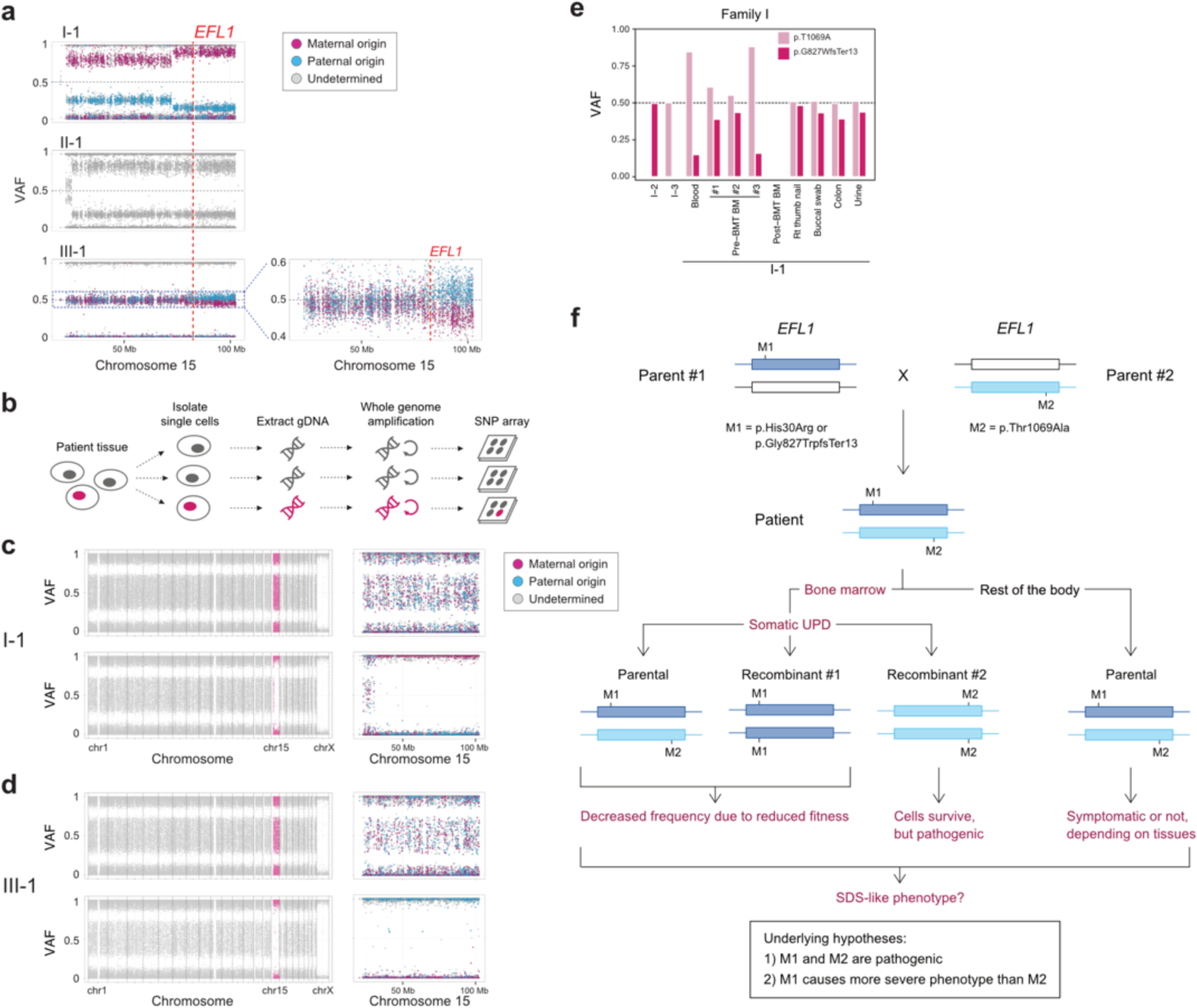
*EFL1* variants display somatic mosaicism. **a** All three patients demonstrate a partial LOH in chromosome 15, as indicated by the deviation from VAF 0.5. The variant origin is indicated by the red or blue dots, whereas parental samples were not available for II-1. The location of *EFL1* is indicated by a red dotted line. **b** Schematic diagram of the single cell LOH experiment shown in c-d, where a red cell symbolizes an LOH-carrying cell. **c-d** Single cell LOH profiles from I-1 bone marrow (**c**) and III-1 buccal swab samples (**d**), the upper plots denote cells without LOH, and the lower plots represent cells with a complete LOH in chr15. The plots on the right show chr15 with the variant origin shown in red or blue dots. **e** VAF of the *EFL1* variant in multiple tissue samples from I-1. VAF, variant allele frequency. **f** Genetic process underlying selection of *EFL1*^p.Thr1069Ala^ cells in the patients.

### EFL1 deficiency impairs 80S ribosome assembly

Our interpretation of the genetic analysis assumes that the *EFL1* variants are pathogenic, and harbor a gradient of severity (Fig. 2f). More specifically, p.His30Arg possesses a comparable severity with the frameshift variant (p.Gly827TrpfsTer13), and these two are more severe than p.Thr1069Ala. To test this, we measured ribosome assembly function in the presence of the *EFL1* variants, as EFL1 is known to mediate GTP hydrolysis-coupled release of eukaryotic translation initiation factor 6 (eIF6) together with SBDS during the maturation of the 60S ribosome subunit^7,15,16^. Thus, we monitored the ribosomal assembly status of the wild type, siRNA-, or CRISPR/Cas9-mediated ablation of *EFL1* (*EFL1*^*KD*^ or *EFL1*^*−/−*^) in HeLa and K562 cell lines to further elucidate the molecular function of the mutant protein (Fig. 3a; Fig. S12). Polysome profiling of the *EFL1*^*KD*^ and *EFL1*^*−/−*^ cells showed a significantly reduced 80S peak (Fig. 3b-c; Fig. S12). This abnormal polysome profile was completely rescued after the introduction of FLAG-tagged wild type *EFL1* but not by the clones that harbored the mutations (Fig. 3b-c; Fig. S13). Interestingly, *EFL1*^*p.His30Arg*^ failed to rescue the mutant phenotype, whereas *EFL1*^*p.Thr1069Ala*^ displayed a moderate effect on ribosome assembly. These results indicated that EFL1 plays a crucial role in ribosome assembly, and *EFL1*^*p.His30Arg*^ possesses a null function, while *EFL1*^*p.Thr1069Ala*^ is hypomorphic.

**Fig. 3.**
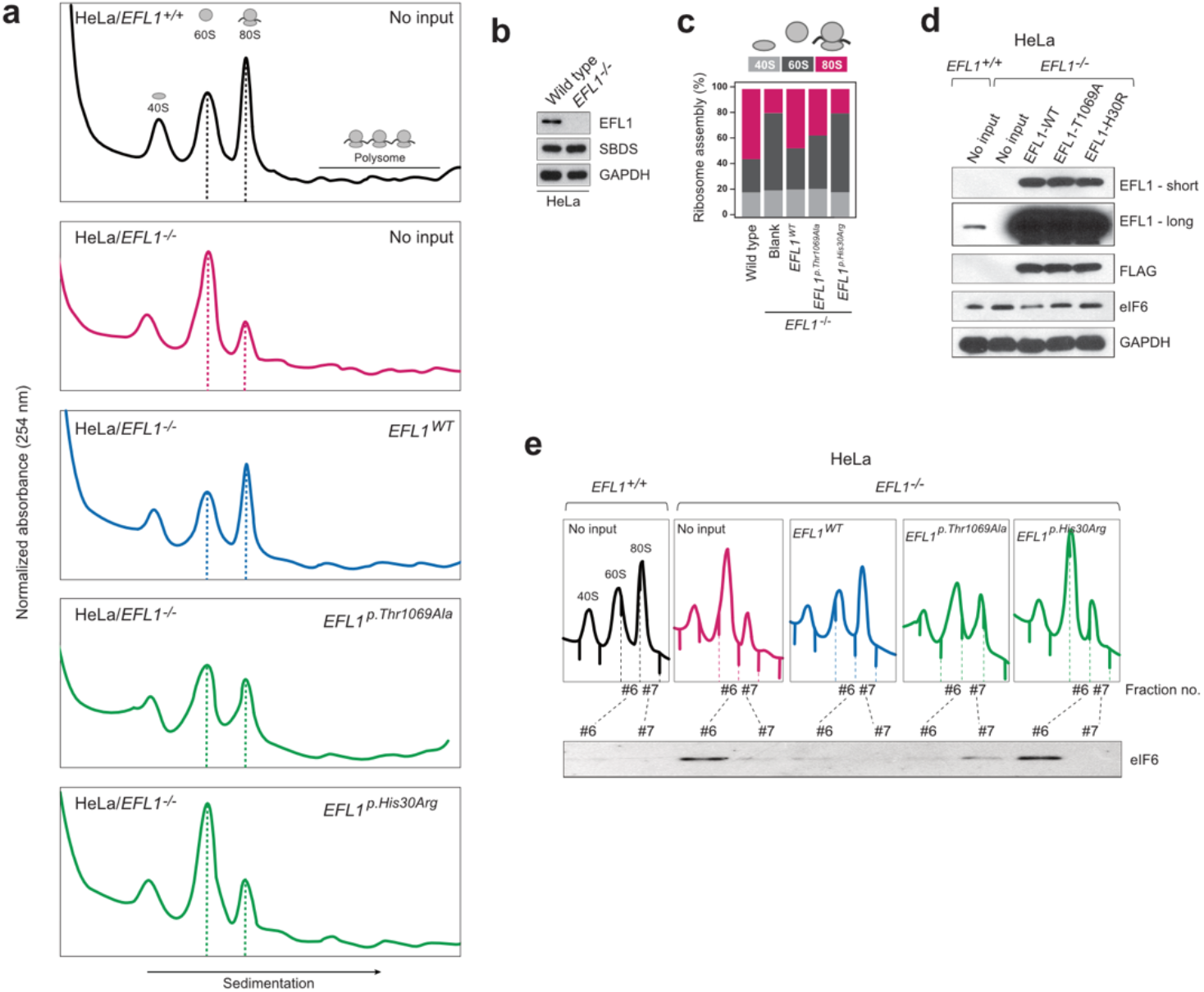
Nonsynonymous variants in *EFL1* disrupt 80S assembly with varying severities. **a** Immunoblot showing the CRISPR-knockout of *EFL1* in HeLa cells. **b** Polysome profiling results of *EFL1*^*+/+*^ and *EFL1*^*−/−*^ cells followed by transfection of vectors containing *EFL1*^*WT*^, *EFL1*^*p.Thr1069Ala*^ or *EFL1*^*p.His30Arg*^ (as indicated in the top right corner of each plot). Note the different heights of 60S and 80S peaks in each experiment (dotted lines). **c** Quantification of the polysome profiling results, displaying the relative occupancy of different ribosome statuses. Each bar corresponds to each experiment in (**b**). **d** Immunoblots of EFL1 and eIF6 in wild type or *EFL1*^−/−^ HeLa cells with or without transfection of vectors containing *EFL1*^*WT*^, *EFL1*^p.Thr1069Ala^ or *EFL1*^p.His30Arg^. **e** Immunoblots of eIF6 from ribosomal fractions that best represent 60S and 80S of HeLa cells, and each corresponds to the polysome profiles shown above.

### Molecular mechanism of EFL1-mediated SDS pathogenesis

Next, we explored the molecular function of the variants in ribosome assembly. Variant function was not mediated via phosphorylation of Thr1069, aberrant subcellular localization of EFL1, or changes in binding affinity to SBDS (Fig. S14). Next, since the release of eIF6 from 60S is a crucial step for 80S assembly and is mediated by SBDS-EFL1, we investigated eIF6 level changes by altering *EFL1*. The assessment of eIF6 in wild type and *EFL1*^−/−^ cells revealed that the absence of *EFL1* induced eIF6 levels, which was partially rescued by the introduction of *EFL1*^*p.Thr1069Ala*^ or *EFL1*^*p.His30Arg*^ (Fig. 3d). Immunoblot analysis of each ribosomal subunit-bound fraction revealed that eIF6 was more highly enriched in 60S ribosome fraction of the *EFL1*^*−/−*^ cells as compared to the wild type or *EFL1*-overexpressed cells (Fig. 3e). Remarkably, introduction of *EFL1*^*p.Thr1069Ala*^ rescued the increased eIF6 in the 60S fraction, whereas *EFL1*^*p.His30Arg*^ failed to do so (Fig. 3e; Fig. S15). Also, absence of *EFL1* caused an impaired shuttling of cytoplasmic eIF6 back to the nucleus, consistent with the previous observation (Fig. S16)^15^. This result, along with the observation that changes in eIF6 and SBDS were not due to transcription levels (Fig. S17), implies that the blocked exclusion of eIF6 and SBDS from the 60S ribosomal subunit is one of the mechanisms by which our mutant protein functions, which results in impaired 80S ribosome assembly. This result also supports our hypothesis that the severity of p.His30Arg is higher than that of p.Thr1069Ala.

### Deficiency of *EFL1* orthologs reproduces SDS phenotypes in zebrafish and mouse models

To determine whether the milder allele (*EFL1*^*p.Thr1069Ala*^) was still pathogenic enough to cause SDS, a morpholino-targeting zebrafish model of *efl1* was subjected to rescue experiments (Fig. S18). The *efl1* morphants had smaller heads and eyes as wells as slightly bent tails and displayed an increased number of apoptotic cells during development (Fig. 4a). Also, primitive erythrocytes and granulocytes were significantly reduced in the *efl1* morphants, indicating impaired primitive hematopoiesis in these embryos (Fig. 4a-b). All phenotypes were rescued by the introduction of the wild type human *EFL1* mRNA, but less so by the *EFL1*^*p.Thr1069Ala*^ mRNA (Fig. 4c-e), confirming that the milder *efl1* allele caused SDS-like features in zebrafish.

**Fig. 4.**
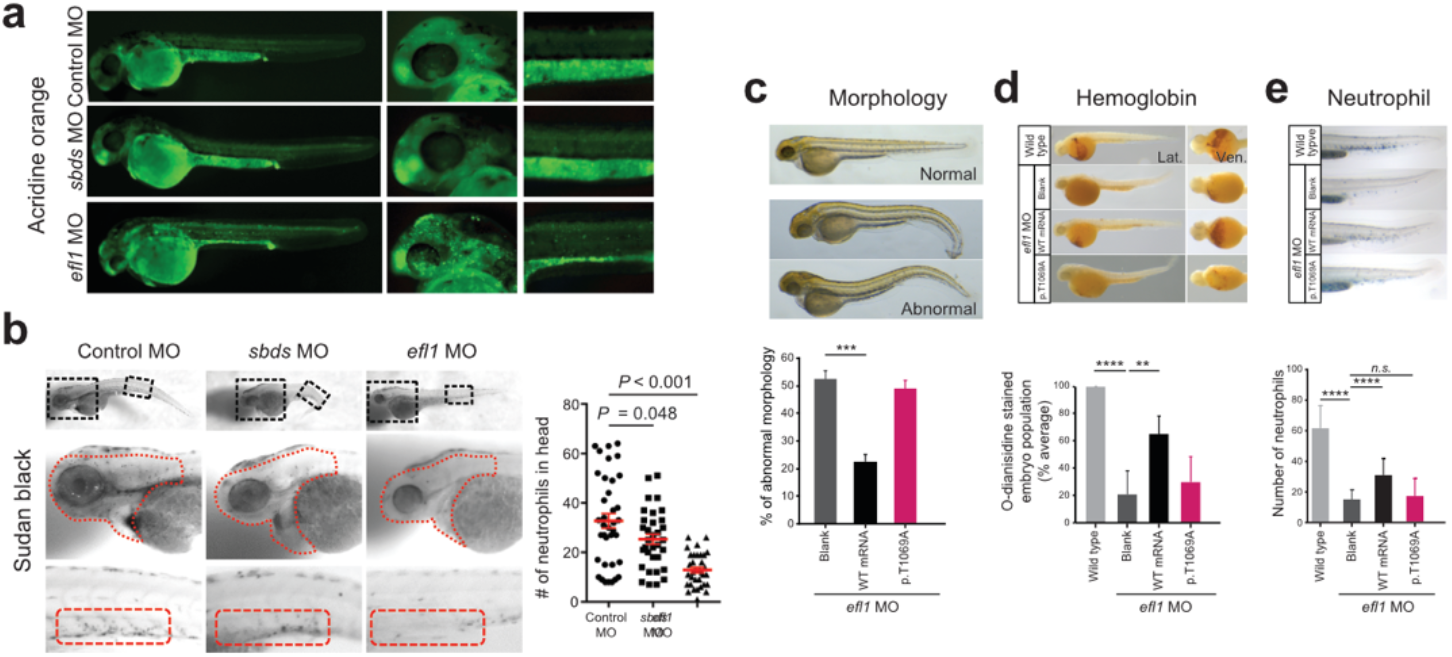
Zebrafish *efl1* mutants model human SDS features. **a-b** Cell death (**a**) and neutrophil production (**b**) of 48 hpf zebrafish embryos by the knockdown of *efl1* and *sbds*. (**c-e**) Rescue of *efl1* MO animals using human *EFL1*^*WT*^ or *EFL1*^*p.Thr1069Ala*^ RNA, documenting morphology (**c**), hemoglobin production (**d**) and neutrophil production (**e**). **, *P* < 0.01; ***, *P* < 0.0005; ****, *P* < 0.0001. hpf, hours post fertilization.

In addition, mouse models were created to further investigate the impact of *Efl1* dysfunction. *Efl1* knock-out and p.Thr1076Ala knock-in mice (mouse Thr1076 is orthologous to human Thr1069; herein designated as *Efl1*^*m*^) were generated and subjected to phenotypic analyses (Fig. S19). Embryos that were homozygous for the null allele (*Efl1*^*−/−*^) were not retrieved on embryonic day (E) 8.5, implying the essential requirement of the gene in early embryogenesis (Fig. S19). On the other hand, mice homozygous for the knock-in allele (*Efl1*^*m/m*^) were viable and healthy (Fig. S19), indicating a differential phenotypic tolerance between mice and humans. To model an accurate *Efl1* dose that may induce an SDS-like phenotypic expression, we inter-crossed the two strains and compared phenotypes of the compound heterozygous (CH) animals (*Efl1*^*m/−*^) to the littermates (*Efl1*^*+/+*^, *Efl1*^*+/−*^ and *Efl1*^*+/m*^), with an emphasis on the major SDS symptoms. The CH mice were smaller (Fig. 5a-b) and died earlier (Fig. 5c). The blood counts revealed reduced hemoglobin, white blood cells and platelets (Fig. 5d), and the bone marrow images displayed a consistent deficiency (Fig. 5e). These results validated our observation that the reduced function of the *EFL1*^*p.Thr1069Ala*^ variant caused SDS in human patients. To determine whether Efl1 expression was affected by the *Efl1*^*m*^ variant, we measured the protein expression from E17.5 livers (Fig. 5f). The liver heterozygous for *Efl1*^*m*^ (*Efl1*^*+/m*^) showed reduced Efl1 expression, suggesting that the variant not only reduced Efl1 activity, but also destabilized the protein. The expression of eIF6 was also detected in the CH livers and was increased compared to the wild type livers, which was consistent with our previous observation in the HeLa cells (Fig. 5g). Ribosome profiling of the wild type and CH livers revealed that the CH livers showed a lower 80S peak compared to the wild type livers (Fig. 5h-i).

**Fig. 5.**
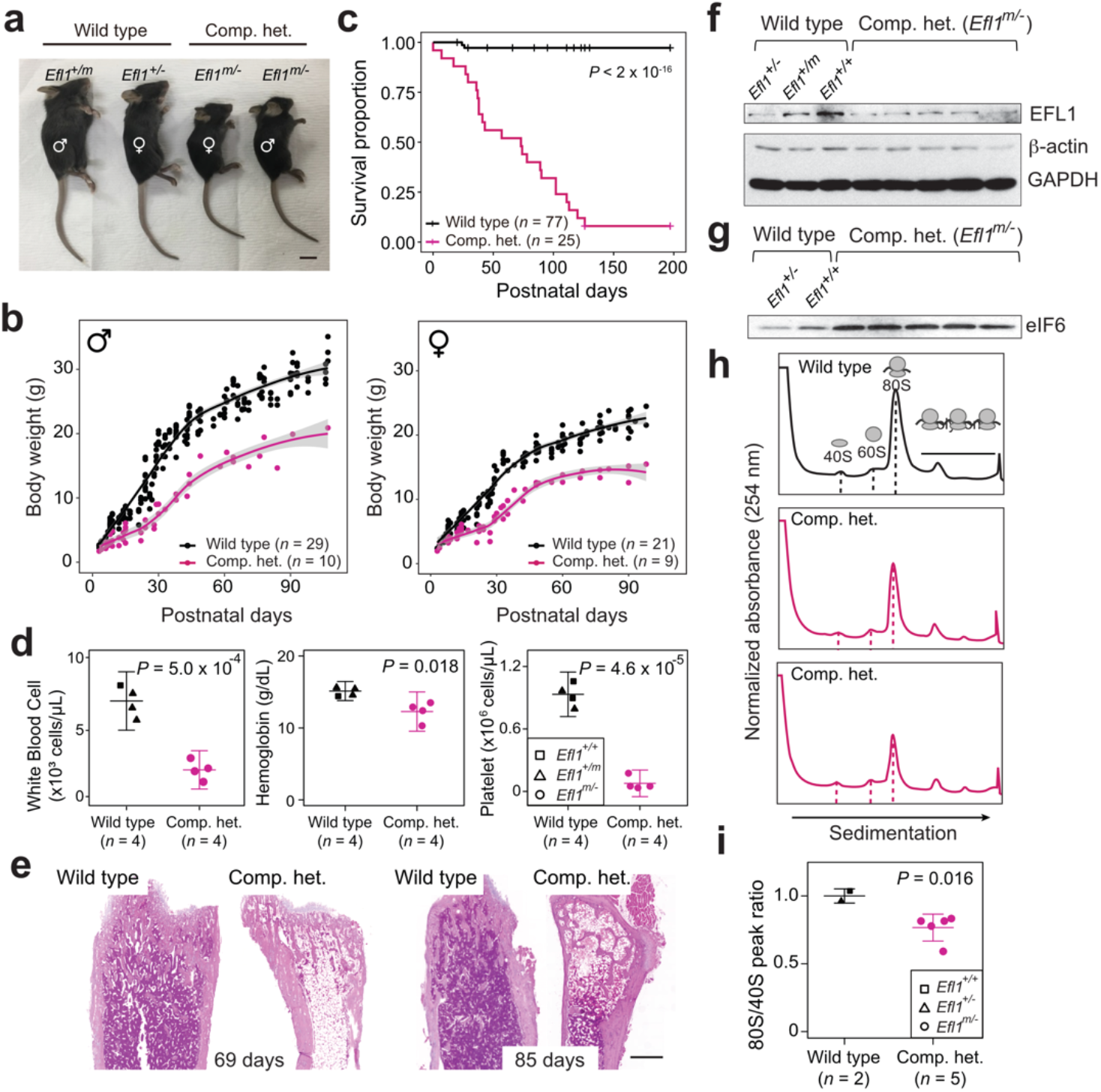
Mouse *Efl1* mutants model human SDS features. **a** Body sizes of the CH mice (*Efl1*^*m/−*^) at three weeks compared to the wild type littermates. **b** Body weight curves of the CH animals compared to the wild types. Males (left) and females (right) are plotted separately. **c** Survival plot. **d** Hematopoietic features of the CH animals, measured at postnatal days 35-42. **e** H&E staining of the bone marrows from 69- and 85-day-old femurs. **f** Level of Efl1 protein in the wild type or CH livers. **g** Immunoblots of eIF6 from wild type or CH livers. **h** Polysome profiling results of the wild type or CH mouse livers. **i** Quantification of the polysome profiling results. Scale bars in (**a**) and (**e**) = 1 cm and 0.5 mm.

### Screening downstream factors of EFL1 dysfunction

To further investigate the features of downstream genes that are strongly affected by the reduced EFL1 function, we performed RNA-seq on the total RNA, 40S-, 80S- and polysome-bound RNAs from wild type and *EFL1*^−/−^ K562 cells (Fig. 6a; Fig. S20). The 60S-bound RNAs were not analyzed as it is unlikely that the large subunit will bind with RNA by itself. The 248 genes that were decreased in the 80S-bound fraction compared to the 40S-bound fraction in the mutant cells were extracted. A gene ontology analysis suggests that RP genes (GO:0003735) were the major constituents of the genes that were reduced by the absence of *EFL1* (*P*_*adj*_ = 5.2 x 10^−77^, adjusted by the Benjamini-Hochberg method; Fig. 6b-c). There was no significantly enriched gene group in the increased gene set. The fractions of the transcripts bound by 40S-, 80S-, or polysome differed substantially for the RP genes in the absence of *EFL1* (*P* < 1.0 x 10^−13^ for differences in both 40S- and 80S-bound transcripts, Wilcoxon signed-rank test), whereas the *TP53* target genes and all other genes did not show such a change (Fig. 6d, *P* > 0.05, Wilcoxon signed-rank test). Next, we compared whether transcript sizes (Fig. S21), expression levels, or consensus sequence elements in the 5’ UTRs may serve as a factor that enabled RP-specific regulation. Notably, highly-expressed RP transcripts with a terminal oligo-pyrimidine (TOP) element (5’-CUUYCUUUUNS-3’) were specifically altered (Fig. 6e; Fig. S22; *P* = 6.9 x 10^−7^). This result revealed that, among all the genes in the genome, 80S ribosome assembly of RP transcripts containing a TOP element were heavily dependent on the normal EFL1 function.

**Fig. 6.**
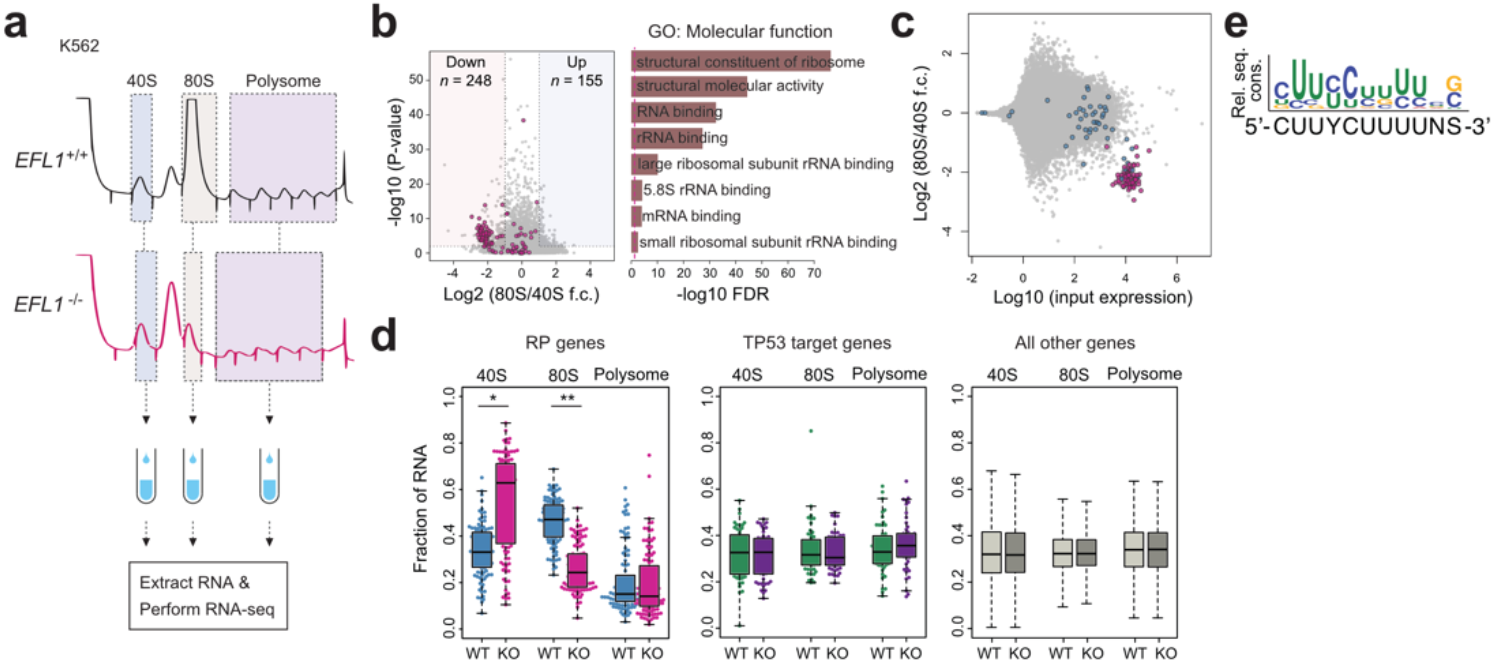
EFL1 is required for RP synthesis. **a** Schematic diagram of the RNA expression analysis from K562 ribosomal fractions. **b** Volcano plot of the genes that were differentially enriched in the 80S fraction compared to the 40S fraction (left). GO analysis of the 248 downregulated genes (right). Red dots depict RP genes (GO:0022625 and GO:0022627, *n* = 118). **c** Fold change of 80S enriched genes normalized by 40S, depicted against transcriptome. Red; significantly downregulated RP genes in (**b**). Blue; other RP genes. **d** Fractions of 40S-, 80S- or polysome-bound RNAs for RP genes (*n* = 108, left), *TP53* target genes (*n* = 51, middle) and all other genes (*n* = 11,812, right) are displayed. Wilcoxon signed-rank test, *. *P* = 4.7 x 10^−14^, **. *P* = 1.5 x 10^−26^. All other test *P*-values > 0.05. **e** Consensus sequence profile of downregulated RP gene 5’ UTRs, deducing a TOP sequence. Rel. seq. cons. denotes relative sequence conservation from the MEME run.

## Discussion

We identified and described a unique bone marrow-specific somatic UPD event that preferentially selects cells with *EFL1* alleles of a weaker severity. Although other parts of the body may suffer from biallelic variants in *EFL1*, composed of more damaging one and the milder one, the hematological system was partially rescued by a somatic UPD resulting in a homozygous *EFL1*^*p.Thr1069Ala*^ with a weaker severity. We provided evidence suggesting that the majority of the cells that carry *EFL1*^*p.Thr1069Ala*^ in a homozygous manner is still pathogenic. We hypothesize that a somatic UPD in the patients could occur and be detected because of the dynamic nature of bone marrow, where the whole stem cell can be potentially replaced by a few clones. Indeed, the somatic UPD was not detected in other solid organs or tissues that were available for investigation. Two lines of evidence suggest infrequent LOH events in the *EFL1* locus in normal populations (2.7-7.7 x 10^−4^; see Results). The odds of having all the p.Thr1069Ala and p.His30Arg variants and the LOH by UPD in the *EFL1* locus for one person by chance is roughly estimated as 1 in 3 trillion, indicating that detectable UPD is not an independent event but is associated with the biallelic *EFL1* variants. It is still not clear if the EFL1 dysfunction somehow contributed to the occurrence of UPD, causing the resulting clone to be expanded by positive selection. Nevertheless, the degree of mosaicism did not seem to directly determine clinical severity (Fig. S11). One could further delineate a more accurate temporal and spatial occurrence of the event if additional clinical samples become available.

Several lines of evidence from our experiments suggest that although p.Thr1069Ala is the mildest among the variants that we found, it is still functional and hypomorphic: (1) the parents of patients that were heterozygous for *EFL1*^*p.Thr1069Ala*^ were asymptomatic, whereas the patients carried the variant in a homozygous status in their bone marrow and showed the pathology; (2) a ribosome profiling assay using a variant allele partially rescued the 80S assembly problem, whereas the wild type allele completely rescued it (Fig. 3); (3) *efl1* MO-treated zebrafish were partially rescued by the variant-containing RNA (Fig. 4); and (4) *Efl1*^*m/−*^ mice displayed an SDS-like phenotype, which was of an intermediate severity relative to the *Efl1*^*−/−*^ and *Efl1*^*+/−*^ mice (Fig. 5). It is notable that although our mouse model successfully phenocopied most of the SDS features, the genotype was not in complete concordance with that of human patients. Our mouse model with *Efl1*^*m/m*^ did not show significant phenotypes, whereas human patients with bone marrow specific homozygous *EFL1*^*p.Thr1069Ala*^ by somatic UPD still had hematological abnormalities such as anemia and neutropenia. This discrepancy between the two species is not rare^23,24^, and it again underscores differences in tolerance to a given variant, perhaps due to different physiological and genetic systems, which is critical in modeling human clinical features in mice.

Nonetheless, we demonstrated that the altered function of EFL1 specifically influenced the translation of RP genes containing a TOP element in the 5’ UTR, which led to a mechanistic mimicry of Diamond-Blackfan anemia, which is another ribosomopathy caused by insufficient RP doses^25,26^. The activation of TP53 is considered as a targetable downstream pathway that leads to SDS or a DBA phenotype^27–29^. However, our data suggest that the loss of *EFL1* does not induce TP53 activation, which is consistent with previous studies of the zebrafish *sbds* model and DBA (Fig. 6d)^30,31^.

It is known that LARP1 directly binds to the TOP element of RP genes to repress translation in a phosphorylation-dependent manner and that mTOR partially regulates LARP1 phosphorylation^32,33^. To determine if we could utilize this pathway to de-repress RP translation and rescue the SDS phenotype in the animal models, we considered a molecular signaling pathway that may regulate RP gene translation through the TOP element. However, both LARP1 binding to RP TOP elements and mTOR signaling were unchanged in the mutant cells, suggesting an alternative mechanism that may regulate RP translation (data not shown).

Here, we demonstrated a mechanism by which biallelic variants of *EFL1* phenocopied classical SDS in three unrelated patients. The bone marrow-specific somatic UPD in these patients mitigated the potentially catastrophic hematological phenotype by homozygosing the less damaging variant (*EFL1*^*p.Thr1069Ala*^). We demonstrated that defective EFL1 caused impaired 80S ribosomal assembly and that the zebrafish and mouse models displayed similar features to humans through the alteration of 80S ribosome assembly of RP transcripts. An extensive search of such SDS patients may provide more insight into the development of somatic mosaicism and subsequent molecular cascades that may lead to new avenues of treatment for ribosomopathy.

## Materials and methods

### Patient recruitment and sampling

Patient enrollment and sampling were conducted under the approval of the Institutional Review Board of Seoul National University Hospital (IRB number: H-1408-014-599). Patients or their parents were provided an informed consent for genetic testing and collecting blood, buccal swab, scalp hair, urine and clipped nails. Biopsy samples of I-1 (esophagus, stomach, duodenum, colon and liver) were retrieved from the Department of Pathology of Seoul National University for a research purpose.

### Whole exome sequencing and variant calling

Trio whole exome sequencing (WES) was performed on one family (I-1, I-2, and I-3) and singleton WES for two families (II-1 and III-1) at Theragen Etex (Suwon, Korea) using genomic DNA extracted from whole blood. Exome was captured using SeqCap EZ Exome v2 Kit (Roche Sequencing, Madison, WI) or SureSelect Human All Exon V5 (Agilent Technologies, Santa Clara, CA) and sequenced by HiSeq 2500 or HiSeq 4000 (Illumina, Inc., San Diego, CA). Paired-end sequencing was performed with read lengths of 75 or 100 base pairs. The raw reads were aligned by BWA MEM software ^34^. Variants were called by Samtools and annotated by in-house pipeline and SnpEff ^34–36^.

### Single nucleus SNP microarray

Nuclei of frozen bone marrow cells from I-1, where a majority of the cells were disrupted during freezing, were prepared and collected manually with a pipette under a phase-contrast microscope. Buccal swab of III-1 was resuspended in 1 ml of PBS and centrifuged at 300 *g* for 3 min. Supernatant was discarded and pellet was resuspended in PBS. This suspension was moved to a 100 ml cell culture dish. Single cells were manually picked up with a pipette under a phase-contrast microscope. These single cells or nuclei were whole-genome amplified by a REPLI-g Single Cell Kit (Qiagen, Venlo, the Netherlands) for 3 hours. Amplified genomes were genotyped by Infinium^®^ OmniExpress-24 v1.2 (Illumina, Inc.).

### Sample-barcoded amplicon sequencing

Genomic DNAs extracted from multiple individuals and tissues were PCR amplified with primers shown in Table S6, with following condition: 3 min at 95°C, followed by 20 cycles (30 sec at 95°C, 30 sec at 57°C, and 20 sec at 72° C), and a final extension at 5 min at 72° C. Barcoded PCRs were performed with 1 µl of first PCR product as templates using tissue-specific barcoded primers (Table S7). Same reverse, forward and reverse primers of *EFL1*, *WDR76* and rs1044032 with Sanger sequencing were used for barcoded PCR products, respectively. The second PCR condition was: 3 min at 95°C, followed by 20 cycles (30 sec at 95°C, 30 sec at 57°C, and 20 sec at 72°C), and a final extension at 5 min at 72°C.

### Molecular dynamics (MD) simulation of EFL1

The initial atom coordinates for MD simulations were from the Protein Data Bank accession number 5ANC ^7^. Input files were prepared with PSFGEN, SOLVATE, and IONIZE plug-ins of VMD ^37^. Each simulation used explicit TIP3 solvent and periodic boundaries. Each unit cell contained one molecule of the entire preinitiation complex. The unit cells were ~210 Å x 180 Å x 190 Å in size and contained ~700,000 atoms. Sodium and chloride were added to make the total ionic strength 100 mM. MD simulations were performed in triplicate (each 50 ns in length) with a CHARMM36 force field ^38^ using NAMD2 ^39^ on the Bebop cluster of the Laboratory Computing Resource Center at Argonne National Laboratory. MD simulations were run at P = 1 atm, T = 310 K, with a 1 fs integration time and a 12 Å cutoff distance. Nonbonded forces were calculated every two steps and electrostatic forces were calculated every four. Pressure and temperature were maintained using a Langevin piston and bath. Atom coordinates were recorded every 10 ps. MD trajectories were visualized and analyzed with VMD and PyMol.

### Polysome profiling

To maintain the binding of mRNA to ribosome subunits, cycloheximide (Sigma-Aldrich, St. Louis, MO) (100 ug/ml) was added to cell culture media and incubated for 10 min at 37°C. After incubation, cells were washed with cold PBS including cycloheximide (10 ug/ml) twice, then lysed with 1 ml of polysome lysis buffer (20 mM HEPES pH 7.6, 5 mM MgCl_2_, 125 mM KCl, 1% NP-40, 2 mM DTT) supplemented with cycloheximide (100 ug/ml), protease inhibitor cocktail (EDTA-free; Roche) and RNase inhibitor (Invitrogen, Carlsbad, CA) on ice. Cell lysates were tumbled for 20 min at 4°C and centrifuged at 13,200 rpm for 20 min. The supernatants were fractionated in 17.5-50% linear sucrose gradients by ultracentrifugation (35,000 rpm for 160 min) in a Beckman ultracentrifuge using SW41-Ti rotor. Gradients were eluted with a gradient fractionator (Brandel, Gaithersburg, MD) and monitored with a UA-5 detector (ISCO). Equal volume of each polysome fraction was used for determining the level of eIF6 by Western blot analysis.

### RNA sequencing from polysome fractions

RNA was extracted from polysome fractionation. RNA sequencing library was constructed using a TruSeq stranded total RNA kit (Illumina Inc.) with rRNA depletion. 100 bp paired-end sequencing was performed using HiSeq 2500, producing >5 Gb for each sample. Transcripts with polysome-bound RNA count >10 were selected and subjected to calculation of fractions of 40S-, 80S- or polysome-bound RNA molecules for a given gene.

### Zebrafish experiments

Zebrafish embryos were co-injected with *efl1* morpholino (MO) and normal or mutated human *EFL1* mRNA at 1-2 cell stage. To evaluate the rescue efficacy of each co-injected *EFL1* mRNA, the number of neutrophils and primary erythrocytes were assessed. For neutrophils, individual Sudan Black stained cells in the caudal region of the embryo, posterior to the anal opening, were counted and the absolute number of positive cells were used for phenotypic comparison. For primary erythrocytes, the degree of o-dianisidine staining was qualitatively compared using the percentage of o-dianisidine stained embryos within the population, since it is not technically possible to count individual erythrocytes.

### Efl1 mutant mouse strain construction, maintenance and experiments

All the mouse experiments were performed under the standard protocols approved by IACUC (#17-0148-S1A0). *Efl1* knock-out (*Efl1*^*−*^) strain was constructed by introducing a 10-base deletion in the 10^th^ exon of the gene in C57BL6/J strain using CRISPR/Cas9 system in Macrogen (Seoul, Korea). One cell embryos were microinjected with two sgRNAs (5’-ACTTCTTTAGGATTAAAAATTGG-3’ and 5’-CCGAGGACAGCGTGGGATATGGG-3’) and Cas9 protein mixture, incubated and transplanted into pseudopregnant recipient ICR mice. *Efl1* knock-in (*Efl1*^*p.Thr1076Ala*^) variant was generated in C57BL6/J in University of Utah Mutation Generation and Analysis Core using two sgRNAs (5’-GTTCTGGGTGCCGACCACGG-3’ and 5’-GTGCAGGTACTCCTCCTCCG-3’) and in the presence of oligodeoxynucleotides which includes the ACC>GCC change mimicking the p.Thr1069Ala and an *Mbo*I site for genotyping. Genotyping strategies of the wild type and mutant alleles are described in supplemental Methods. For phenotypic evaluations, wild type and mutant mice from the same litter were sacrificed and dissected for peripheral blood extraction, pancreas pathology and skeletal structure analysis. 300 ul of peripheral blood was extracted from 35-42-day animals and subjected to a complete blood count and differential tests.

## Author contributions

M. Choi, H.J. Kang, and H.H. Kim conceived the study. S. Lee and M. Choi performed genetic analysis and statistical evaluations. C.H. Shin, S. Lee, S.D. Jeong, H.H. Kim, J. Lee, and M. Choi performed molecular biology and biochemistry experiments and assessed the results. J. Lee, S. Lee, S.J. Son, M. Choi, and J.K. Seong performed mouse experiments. J.-D. Kim, A.-R. Kim, and S.-W. Jin performed zebrafish experiments. C.R. Hong, J.S. Ko, Y.B. Sohn, O.-H. Kim, J.M. Ko, T.-J. Cho, and H.J. Kang provided patient care and generated clinical data. N.T. Wright, O. Kokhan, and T. Yoo analyzed protein structure. M. Choi, S. Lee, C.H. Shin, C.R. Hong, J. Lee, H.J. Kang, and H.H. Kim wrote the manuscript. All authors approved the final version of the manuscript.

## Acknowledgements

We appreciate the patients and families that participated in this study. We thank Jung-Ah Kim and Hyoung-Jin Kim at Seoul National University Hospital for collecting and interpreting hematological data and for interpreting images. We thank the University of Utah Mutation Generation and Detection Core for construction of the *Efl1* knock-in mice. This study was partly supported by grants from the National Research Foundation of Korea (2014M3C9A2064686, 2019R1A2C2010789 to M.C. and 2017R1A2A2A05069691 to H.H.K.) and National Science Foundation (REU CHE-1062629 and RUI MCB-1607024) awards to N.T.W.. This research was partly supported by Korea Mouse Phenotyping Project (2013M3A9D5072550) of the Ministry of Science, ICT and Future Planning through the National Research Foundation. OK gratefully acknowledges the computing resources provided by Bebop, a high-performance computing cluster operated by the Laboratory Computing Resource Center at Argonne National Laboratory. The authors declare no competing financial interests.

## Abbreviations used

CH: compound heterozygous
DBA: Diamond-Blackfan anemia
LOH: loss of heterozygosity
RP: ribosomal protein
SDS: Shwachman-Diamond syndrome
TOP: terminal oligo-pyrimidine
UPD: uniparental disomy
VUS: variants of unknown significances

## Notes

### Competing Interest Statement

The authors have declared no competing interest.

