## Supplementary materials for "Somatic uniparental disomy mitigates the most damaging *EFL1* allele combination in Shwachman-Diamond syndrome"

### Contents


| | |
|---|---|
| Methods | 2 |
| Clinical Narratives | 8 |
| Supplemental Tables | 10 |
| Supplemental Figures | 19 |
| References | 42 |

**Methods**

*Sanger sequencing*

PCR amplification was performed with 5 pmol of each specific primer (Supplemental Table 4). The PCR conditions were: 95°C for 3 min, followed by 35°C cycles (95°C for 30 s, 57°C for 30 s, and 72°C for 20 s), and a final extension at 72°C for 5 min. Sanger sequencing reactions were run on an ABI3730XL DNA Analyzer (Applied Biosystems, Waltham, MA).

*Orthologs*

Proteins encoded by orthologs of EFL1 in multiple species were identified by a BLAST search with the human sequence as bait (NP_078856.4). GenBank accessions for these are NP_780526.2 (*Mus musculus*), XP_002721510.1 (*Oryctolagus cuniculus*), XP_002696652.1 (*Bos taurus*), NP_001034345.2 (*Gallus gallus*), NP_001084970.1 (*Xenopus laevis*), XP_697184.5 (*Danio rerio*), XP_002120234.2 (*Ciona intestinalis*) and NP_788515.1 (*Drosophila melanogaster*). Protein sequences were aligned by ClustalW2 software (Larkin et al., 2007).

*SNP microarray*

SNP genotyping was performed on Infinium® OmniExpress-24 v1.1 BeadChip (Illumina, Inc.) or Infinium® OmniExpress-24 v1.2 (Illumina, Inc.) at Yale Center for Genome Analysis and raw data were extracted and analyzed using GenomeStudio (Illumina, Inc.).

*CRISPR/Cas9 system design*

Expression vector with a *Cas9* gene (pRGEN-Cas9-CMV) and single guide RNA (sgRNA) plasmid were prepared from ToolGen, Inc. (Seoul, Korea). Two sgRNAs were designed to target exons 8 (5'- TAT ATA GTA ATC TCC CCA CAA GG-3')
and 10 (5'-GGT CTG AAT GTC GTG CCT CCC GG-3') of *EFL1*, of which the last 3 nucleotides were PAM sequences.

*Cell culture*

HeLa cell line was maintained in Dulbecco's Modified Eagle's medium (DMEM; HyClone™, GE Healthcare Life Sciences, Chicago, IL) supplemented with 10% Fetal Bovine Serum (FBS; HyClone™, GE Healthcare Life Sciences) and 1% penicillin/streptomycin (WELGENE, Gyeongsan, Korea). K562 cell line was maintained in the Roswell Park Memorial Institute 1640 medium (RPMI 1640; HyClone™, GE Healthcare Life Sciences) supplemented with 10% Fetal Bovine Serum (FBS; HyClone™, GE Healthcare Life Sciences) and 1% penicillin/streptomycin (WELGENE).

*EFL1 knockout in cell lines by CRISPR/Cas9*

*EFL1* knockout in HeLa cells was performed by transfection of two plasmids expressing Cas9 and sgRNA. Briefly, HeLa cells were seeded on 6-well cell culture plates a day before transfection. On the transfection day, cells were about 70 % confluent. Culture media was removed and replaced with 700 µl of Opti-MEM (Gibco, Waltham, MA). 1.25 µg of sgRNA plasmid and 1.25 µg of Cas9 plasmid were mixed with 150 µl of Opti-MEM (tube 1). 10 µl of Lipofectamine 2000 (Invitrogen, Waltham, MA) was mixed with 150 µl of Opti-MEM (tube 2). Tube 1 and tube 2 were mixed to make 300 µl of transfection mixture and incubated for 25 min. Transfection mixture was dropped to the media and incubated in 37°C for 6 hours. Then, Opti-MEM was removed, washed once with PBS, and replaced with 2 ml of new media. *EFL1* knockout in K562 cell was generated by transfection of two plasmids expressing Cas9 and sgRNA by electroporation. Briefly, $4 \times 10^5$ / 100 µl concentration of K562 cells in BTX High Performance Electroporation solution were moved to BTX 2 mm electroporation cuvette along with 1 ug of sgRNA and 1 µg of Cas9 plasmids. Electroporation was performed with one pulse of 200 V and 9 ms duration, using BTX ECM 830. Electroporated cells were moved to a 25T flask with new media. Two K562 clones (#6 and #7) were selected, maintained and utilized for subsequent experiments.

*T7E1 assay for assessment of CRISPR/Cas9 activity*

T7E1 assay was done using a EnGen® Mutation Detection Kit (NEB, Ipswich, MA) for screening of CRISPR/Cas9-induced mutation. Genomic DNA was PCR amplified using corresponding primer sets; forward primer: 5'-GAC CTG AAC ATG GGC TTT GG-3' and reverse primer: 5'-TCC CTC ATG AAC TCA CCT GC-3' for sgRNA targeting exon 8 of *EFL1*, and forward primer: 5'-TGG GTT CCA CTG CCT GTT TA-3' and reverse primer: 5'-TGG CTA CAG GTG ACC TTG AA-3' for sgRNA targeting exon 10 of *EFL1*. PCR product was used for downstream heteroduplex formation and following T7 endonuclease digestion according to the protocol provided by the manufacturer.

*Clonal selection of knockout cell line*

After CRISPR/Cas9 activity was validated by a T7E1 assay, clonal selection of cell was performed by limiting dilution. Briefly, cells were counted and diluted to a concentration of 0.5 cells/100 μl and seeded to flat-bottom 96-well cell culture plates with 100 μl/well. Growth of cell colonies was monitored for 3 weeks. Colonies were moved to a larger culture plate after sufficient growth. Sanger sequencing was performed to make sure of knockout alleles of each colony after genomic DNA was PCR amplified using identical primer sets that were used for the T7E1 assay.

*EFL1 plasmid construction and rescue experiment*

*EFL1* cDNA clone was obtained from Addgene (https://www.addgene.org/) and subcloned to pcDNA-3X-FLAG EFL1 WT. For *EFL1* mutant construction (EFL1 p.Thr1069Ala and p.Thr1069Asp), a QuickChange site-directed mutagenesis kit (Stratagene, CA) was used with each primer (Forward: 5'- CCC CTT CTG GGT GCC AGC TAC TGA GGA GGA ATA C-3' and Reverse:5'- GTA TTC CTC CTC AGT AGC TGG CAC CCA GAA GGG G-3' for p.Thr1069Ala and 5'- CCC TTC TGG GTG CCA GAT ACT GAG GAG GAA TAC-3' and 5'-GTA TTC CTC CTC AGT ATC TGG CAC CCA GAA GGG-3' for p.Thr1069Asp) designed in Primer-X according to the manufacturer's protocol. Blank vector and vectors expressing wild-type *EFL1*, p.Thr1069Ala *EFL1*, and p.Thr1069Asp *EFL1* were used for the rescue experiment. CRISPR-mediated *EFL1* knockout HeLa cells were seeded to a 150 mm culture dish a day before transfection. Cells were transfected with vectors using Lipofectamine 2000 (Invitrogen) at ~70% confluency according to the manufacturer's protocol.

*Western blot*

Cells were lysed using radioimmunoprecipitation (RIPA) buffer containing protease and phosphatase inhibitors (Roche, Basel, Switzerland). Equal amounts of cell lysate were separated by SDS–polyacrylamide gel electrophoresis and transferred to polyvinylidene difluoride membranes (Millipore, Billerica, MA, USA). After blocking, membranes were incubated with the respective primary antibody, washed, and incubated with the appropriate secondary antibody (Supplemental Table 6).

*Subcellular fractionation*

For isolation of the cytosolic and nuclear fraction, cells were harvested and lysed with digitonin (Sigma, St. Louis, MO)-containing RSB buffer. After centrifugation at 2,000 *g* for 10 min, the supernatant was transferred to a new tube (Cytosolic extracts). The remaining pellet was washed five times with RSB buffer and lysed with RIPA buffer. Nuclear extracts were isolated by centrifugation at 13,200 rpm for 20 min.

*Gene ontology analysis*

List of genes were submitted to ToppFun of ToppGene Suite and analyzed with default thresholds (Chen et al., 2009).

*Motif enrichment test*

Enriched motif in 5' UTR of genes was searched using MEME (Multiple Em for Motif Elicitation) of MEME Suite (version 5.0.2) (Bailey et al., 2009).

*Zebrafish husbandry and morpholino (MO) and mRNA injection*

Zebrafish (*Danio rerio*) were maintained as previously described (Westerfield, 2000). Control, *efl1* and *sbds* morpholinos (MOs) were purchased from Genetools. MOs were injected with phenol-red (0.1%) in zebrafish embryos at a 1-2 cell stage and the efficacy of MO was evaluated by semi-quantitative conventional RT-PCR before experiment. The sequences of MO used were *efl1* MO; 5'-ATG TTG GAT GAT ACC TGC GGA CAC A-3' and *sbds* MO; 5'-GCT TTG ACC ATT CAG ATC ACC TGT T-3′. mRNA was *in vitro* transcribed using normal or mutated human *EFL1* cDNA as a template and similarly injected at a 1-2 cell stage together with MO.

*Acridine orange staining in zebrafish*

Acridine orange (Sigma) was prepared as a stock solution (300x) of 5 mg/ml in egg water and diluted to 1x concentration in use. Control and morphants were dechorionated at 48 hours-post-fertilization (hpf), and embryos were incubated for 20 min in room temperature and in the dark. Then embryos were washed 3 times in egg water, and imaged using a fluorescence microscope.

*o-Dianisidine staining in zebrafish*

Control, *sbds* and *efl1* MOs were injected at a 1-2 cell stage and collected at 48 hpf. Non-fixed dechorionated embryos were incubated in solution containing o-dianisidinestained (0.6 mg/ml), 0.01 M sodium acetate (pH 4.5), 0.65% $H_2O_2$, and 40% (vol/vol) ethanol for 15 min in the dark. Then, stained embryos were washed with benzyl benzoate/benzyl alcohol (2:1, vol/vol) until the background signal was cleared.

*Sudan black staining in zebrafish*

Control, *sbds* and *efl1* MO-injected embryos were fixed by 4% paraformaldehyde at 48 hpf. Fixed embryos were rinsed 3 times in PBS and stained by Sudan Black solution (Sigma) for

20 min. Stained embryos were washed in 70% ethanol, and images were taken under a microscope. The number of neutrophils was counted in the head and ventral tail regions.

*Mouse genotyping*

To genotype *Efl1* knockout allele, mouse genomic DNA was amplified using a forward primer 5'-GCC TTT TCT CTC TGA TCT TTC CC-3' and reverse primer 5'-GAG GAC AGC GTG GGA TAT GG-3' under the PCR condition of initial denaturation at 94 ºC for 5 min, 35 cycles of 94 ºC 30 sec, 55 ºC 30 sec and 72 ºC 20 sec, and final extension of 72 ºC 20 sec. Wild type and knockout alleles yield 180 and 170 bp products, respectively. To genotype $Efl1^{p.Thr1076Ala}$ knock-in allele, genomic DNA was initially amplified using a forward primer 5'-AAC ATC CAC CTC ACC TAG CC=3' and reverse primer 5'-TTC CCG GAT TAG ACC CTT GA-3' using the PCR condition of initial denaturation at 94 ºC for 5 min, 50 cycles of 94 ºC 30 sec, 56 ºC 30 sec and 72 ºC 20 sec, and final extension of 72 ºC 5 min. The 316 bp product was digested with *Mbo*I (New England Biolabs, Ipswich, MA), to distinguish wild type (non-digestible) and knock-in (cut into 258 and 58 bp fragments) alleles.

### 2. Clinical narratives

*I-1*

Proband I-1 is a 3-year-old boy who had severe intrauterine growth retardation resulting in preterm (35+3 weeks) delivery with a 1.7 kg of birth weight. He had thrombocytopenia on the day of birth, and neutropenia and anemia after 3 weeks and 2 months after birth, respectively. Neonatal metabolic screening tests were normal. Bone marrow examination performed at 6 months of age revealed hypocellularity, reduced megakaryocytes and an increased storage of iron. His hematological profiles did not recover since then. At the age of 1, ultrasonography and magnetic resonance imaging (MRI) revealed pancreatic lipomatosis and liver hemochromatosis (Fig. 1a, upper panel), along with exocrine pancreatic insufficiency which was diagnosed by alpha-1 anti-trypsin (<8.5 mg/dL) and pancreatic elastase (<5.5 ng/mL) levels of stool specimen. He also had generalized osteoporosis with metaphyseal chondrodysplasia (Fig. 1a, lower panel). He underwent successful allogenic umbilical cord hematopoietic stem cell transplantation (allogeneic HSCT), resulting in complete recovery of hematologic profiles. *SBDS* gene sequencing did not show plausible pathogenic variants.

*II-1*

II-1 is a 9-year-old girl who had severe intrauterine growth retardation resulting in preterm (36 weeks) delivery with a 1.6 kg of birth weight. She had thrombocytopenia which was spontaneously resolved at 3 years of age. She was short and showed poor weight gain, and suffered from chronic diarrhea and steatorrhea since early childhood. At an initial survey, upper gastrointestinal and small bowel barium examination was performed to search for the cause of diarrhea. Then her abdominal radiograph revealed the proximal femoral metaphyseal irregularity, and further skeletal survey found narrow thorax and short ribs. Metaphyseal dysplasia was also noted at the knees. Concurrent abdominal CT revealed atrophic pancreas with fatty infiltration (Fig. 1a). She also had a developmental delay. With a clinical history of chronic diarrhea and skeletal abnormality, SBDS was suspected as a causal gene, but no clear pathogenic variant was identified.

*III-1*

III-1 is a 25-year-old male who did not have any perinatal problems except low birth weight (40+4 weeks, 2.4 kg). At the age of 3 months, he had pneumonia and thrombocytopenia, while bone marrow examination did not show any significant abnormalities. At two years old, he had pancreatic exocrine and endocrine insufficiencies, thrombocytopenia, anemia, intermittent neutropenia, metaphyseal chondrodysplasia and ichthyosis which led to a clinical diagnosis of SDS (Figure 1A). On longer follow-up, he was found to have osteoporosis, hepatomegaly and total fatty change of the pancreas (Figure 1A). He had developmental delay and currently has mental retardation. His *SBDS* gene test result was normal.

**Supplemental Table 1.** Clinical information of three patients

| Proband | **I-1** | **II-1** | **III-1** |
|---|---|---|---|
| Sex | Male | Female | Male |
| Gestational age at birth | 35+3 weeks | 36 weeks | 40+4 weeks |
| Birth weight | 1.7 kg | 1.6 kg | 2.4 kg |
| Age of onset | 3 weeks | <3.5 years, presumably after birth | 3 months |
| Pancreas lipomatosis | + | + | + |
| Exocrine pancreatic insufficiency | + | n.a. | + |
| Hematologic manifestation | Anemia Thrombocytopenia Neutropenia | Thrombocytopenia | Anemia Thrombocytopenia, Intermittent neutropenia |
| Metaphyseal chondrodysplasia | + | + | + |
| Osteopenia/osteoporosis | + | + | + |
| Developmental delay | + | + | + |

*. n.a.: not assessed

**Supplemental Table 2.** WES run summary

| | I-1 | I-2 | I-3 | II-1 | III-1 |
|---|---|---|---|---|---|
| Read length (bp) | 2 x 74 | 2 x 74 | 2 x 74 | 2 x 101 | 2 x 101 |
| Number of reads (millions) | 67.5 | 72 | 61.9 | 77.3 | 80.6 |
| Mean coverage depth (X) | 62.7 | 67.3 | 56.9 | 88.1 | 75.7 |
| % of reads on genome | 91.51 | 91.56 | 91.45 | 99.35 | 98.84 |
| % of reads on target | 60.77 | 61.15 | 60.31 | 81.39 | 65.22 |
| % of bases covered at least 4x | 97.44 | 97.63 | 97.32 | 99.63 | 99.63 |
| % of bases covered at least 8x | 95.13 | 95.63 | 94.72 | 99.23 | 99.01 |
| % of bases covered at least 20x | 84.04 | 86.03 | 82.19 | 96.48 | 94.47 |
| % of per-base error rate | 0.41 | 0.40 | 0.39 | 0.34 | 0.37 |
| % of PCR duplicate | 5.79 | 5.97 | 5.20 | 4.70 | 2.53 |

**Supplemental Table 3.** Rare functional variants in chromosome 15 called by WES.

| Patient | Gene | Type | AA change | AA position/ Length | PhyloP | dbSNP ID | 1000G AF | ExAC AF | gnomAD AF | Chr | Position (hg19) | Base change | Ref cov | Nonref cov | Nonref cov/total cov |
|---|---|---|---|---|---|---|---|---|---|---|---|---|---|---|---|
| I-1 | *WDR76* | Missense | p.Ser614Gly | 614/626 | 6.202 | rs3742985 | 4.00E-03 | 1.57E-03 | 1.52E-03 | chr15 | 44158549 | A>G | 9 | 63 | 0.88 |
| I-1 | *C15orf60* | Missense | p.Val101Leu | 101/266 | -1.166 | rs182542888 | 1.00E-03 | 7.04E-04 | 4.08E-04 | chr15 | 73832877 | G>T | 19 | 140 | 0.88 |
| **I-1** | ***EFL1*** | **Missense** | **p.Thr1069Ala** | **1069/1120** | **7.733** | **rs756494164** | **0.00E+00** | **2.48E-05** | **3.26E-05** | **chr15** | **82422872** | **T>C** | **10** | **64** | **0.86** |
| II-1 | *APBA2* | Missense | p.Pro171Leu | 171/752 | 1.252 | rs148760039 | 1.13E-02 | 1.93E-03 | 1.92E-03 | chr15 | 29346590 | C>T | 29 | 119 | 0.80 |
| II-1 | *PLA2G4D* | Missense | p.Arg691Trp | 691/818 | 0.003 | rs766201188 | 0.00E+00 | 3.70E-05 | 2.46E-05 | chr15 | 42362266 | G>A | 15 | 8 | 0.35 |
| II-1 | *MAP1A* | Missense | p.Thr2643Ala | 2643/3041 | 2.901 | rs192770595 | 7.10E-03 | 9.60E-04 | 9.78E-04 | chr15 | 43820884 | A>G | 10 | 50 | 0.83 |
| II-1 | *NEDD4* | Missense | p.Gly479Glu | 479/1319 | 1.929 | rs200370285 | 0.00E+00 | 1.40E-04 | 1.19E-04 | chr15 | 56207594 | C>T | 16 | 45 | 0.74 |
| II-1 | *C2CD4B* | Missense | p.Thr298Ile | 298/364 | 2.783 | rs373533878 | 0.00E+00 | 5.43E-04 | 6.66E-04 | chr15 | 62456291 | G>A | 0 | 62 | 1.00 |
| II-1 | *MYO9A* | Missense | p.Asn1508Ser | 1508/2619 | 4.663 | rs778441953 | 0.00E+00 | 8.24E-05 | 5.69E-05 | chr15 | 72190321 | T>C | 31 | 121 | 0.80 |
| **II-1** | ***EFL1*** | **Missense** | **p.Thr1069Ala** | **1069/1120** | **7.733** | **rs756494164** | **0.00E+00** | **2.48E-05** | **3.26E-05** | **chr15** | **82422872** | **T>C** | **17** | **59** | **0.77** |
| II-1 | *AKAP13* | Missense | p.Thr339Ser | 339/2817 | -1.566 | rs79064356 | 0.00E+00 | 7.91E-04 | 7.69E-04 | chr15 | 86122314 | A>T | 125 | 37 | 0.23 |
| II-1 | *ACAN* | Missense | p.Gly395Ser | 395/2530 | 5.240 | rs117772298 | 0.00E+00 | 1.10E-03 | 1.01E-03 | chr15 | 89388867 | G>A | 124 | 29 | 0.19 |
| III-1 | *C15orf55* | Missense | p.Thr809Met | 809/1160 | -1.669 | rs16959028 | 5.39E-03 | 3.28E-03 | 3.19E-03 | chr15 | 34648635 | C>T | 31 | 32 | 0.51 |
| III-1 | *SPTBN5* | Missense | p.Arg1367Thr | 1367/3674 | 0.147 | rs2290558 | 4.59E-03 | 2.70E-03 | 2.71E-03 | chr15 | 42168334 | C>G | 18 | 14 | 0.44 |
| III-1 | *DUOXA1* | Missense | p.Arg133Leu | 133/483 | -1.295 | rs75981505 | 1.16E-02 | 4.78E-03 | 4.56E-03 | chr15 | 45412946 | C>A | 38 | 28 | 0.42 |
| III-1 | *AQP9* | Missense | p.Val120Ile | 120/295 | 3.715 | rs200107166 | 3.99E-04 | 4.94E-05 | 5.77E-05 | chr15 | 58465386 | G>A | 15 | 10 | 0.40 |
| III-1 | *BNIP2* | Missense | p.Pro53Leu | 53/435 | -1.751 | rs3087331 | 2.60E-03 | 5.10E-04 | 1.49E-03 | chr15 | 59981482 | G>A | 21 | 20 | 0.48 |
| **III-1** | ***EFL1*** | **Missense** | **p.Thr1069Ala** | **1069/1120** | **7.733** | **rs756494164** | **0.00E+00** | **2.48E-05** | **3.26E-05** | **chr15** | **82422872** | **T>C** | **43** | **30** | **0.41** |
| III-1 | *NGRN* | Missense | p.Leu7Pro | 7/291 | 0.039 | rs182240673 | 2.40E-03 | 1.03E-03 | 1.07E-03 | chr15 | 90808964 | T>C | 20 | 9 | 0.31 |
| III-1 | *MAN2A2* | Missense | p.Asp891Gly | 891/1150 | 5.880 | rs75478132 | 4.19E-03 | 1.47E-03 | 1.35E-03 | chr15 | 91456590 | A>G | 20 | 17 | 0.45 |

| | | | | | | | | | | | | | | | |
|---|---|---|---|---|---|---|---|---|---|---|---|---|---|---|---|
| III-1 | *CHD2* | Missense | p.Ala1100Pro | 1100/1828 | 9.300 | rs763614224 | 0.00E+00 | 4.12E-05 | 2.85E-05 | chr15 | 93528788 | G>C | 52 | 40 | 0.43 |
| III-1 | *ARRDC4* | Missense | p.Pro49Ser | 49/418 | 1.145 | rs202148289 | 6.79E-03 | 1.21E-03 | 2.47E-03 | chr15 | 98504236 | C>T | 14 | 10 | 0.42 |

AA; amino acid, AF; allele frequency, cov; coverage

**Supplemental Table 4.** Evaluation of EFL1 p.Thr1069Ala, p.His1069Arg and SBDS p.Asp110Asn using variant functionality evaluation software.

| Variant | Position (hg19) | SIFT | Polyphen2 HVAR | LRT | MutationTaster | CADD | GERP++ NR | phyloP100way |
|---|---|---|---|---|---|---|---|---|
| *EFL1*$^{p.Thr1069Ala}$ | chr15:82130531 | 0.014 | 0.804 | 0.000832 | 1 | 4.11 | 5.18 | 7.75 |
| *EFL1*$^{p.His30Arg}$ | chr15:82261690 | 0.002 | 0.999 | 0.000722 | 1 | 4.21 | 3.82 | 7.05 |
| *SBDS*$^{p.Asp110Asn}$ | chr7:66993348 | 0.039 | 0.148 | 0 | 1 | 3.02 | 4.47 | 6.85 |

**Supplemental Table 5.** List of reported *EFL1* variants from SDS patients.

| Genomic coordinate (hg19) | cDNA (NM_024580.5) | Protein | Reference |
|---|---|---|---|
| chr15:g.82,422,793C>T | c.3284G>A | p.Arg1095Gln | (Stepensky et al., 2017) |
| chr15:g.82,443,887C>T | c.2908G>A | p.Arg970His | (Tan et al., 2019) |
| chr15:g.82,444,148A>C | c.2647T>G | p.Cys883Gly | (Tan et al., 2019) |
| chr15:g.82,444,150A>T | c.2645T>A | p.Met882Lys | (Stepensky et al., 2017) |
| chr15:g.82,444,535G>A | c.2260C>T | p.Arg754* | (Tan et al., 2019) |
| chr15:g.82,512,090A>G | c.1514T>C | p.Phe505Ser | (Tan et al., 2019) |
| chr15:g.82,532,896T>C | c.379A>G | p.Thr127Ala | (Tan et al., 2018) |

**Supplemental Table 6.** PCR primers used for Sanger sequencing.

| Position (hg19) | Locus | Direction | Nucleotide sequences 5'-3' | Product length (bp) |
|---|---|---|---|---|
| chr15:82,422,872 | *EFL1* | Forward | GCT TTT CTG CAT GCT CCA CA | 235 |
| | | Reverse | GGG GAG CTC AGT GAA TAA GGT | |
| chr15:44,158,549 | *WDR76* | Forward | CAG GAA AGA GGG TGC ATT CG | 224 |
| | | Reverse | TAA ACA CAC TAA GGC ACG CC | |

**Supplemental Table 7.** Barcoded forward primers for *EFL1* and *WDR76* amplicon generation of family I.

| Sample | Gene | Nucleotide sequences 5'-3' |
|---|---|---|
| I-1 (blood) | *EFL1* | AGAT**CGATGT**CTGCGTTCATGTACTTCCGG |
| I-2 (blood) | *EFL1* | AGAT**ACAGTG**CTGCGTTCATGTACTTCCGG |
| I-3 (blood) | *EFL1* | AGAT**TGACCA**CTGCGTTCATGTACTTCCGG |
| BM before uHSCT | *EFL1* | AGAT**GCCAAT**CTGCGTTCATGTACTTCCGG |
| BM after uHSCT | *EFL1* | AGAT**CAGATC**CTGCGTTCATGTACTTCCGG |
| Right thumb nail | *EFL1* | AGAT**CTTGTA**CTGCGTTCATGTACTTCCGG |
| Right toe nail | *EFL1* | AGAT**TTAGGC**CTGCGTTCATGTACTTCCGG |
| Buccal swab | *EFL1* | AGAT**TAGCTT**CTGCGTTCATGTACTTCCGG |
| Duodenum | *EFL1* | AGAT**AGTTCC**CTGCGTTCATGTACTTCCGG |
| Colon | *EFL1* | AGAT**ATGTCA**CTGCGTTCATGTACTTCCGG |
| Urine | *EFL1* | AGAT**GTAGAG**CTGCGTTCATGTACTTCCGG |
| I-1 (blood) | *WDR76* | AGAT**CGATGT**CCGATTACTAGGCTCCTTAGAC |
| I-2 (blood) | *WDR76* | AGAT**ACAGTG**CCGATTACTAGGCTCCTTAGAC |
| I-3 (blood) | *WDR76* | AGAT**TGACCA**CCGATTACTAGGCTCCTTAGAC |
| BM before uHSCT | *WDR76* | AGAT**GCCAAT**CCGATTACTAGGCTCCTTAGAC |
| BM after uHSCT | *WDR76* | AGAT**CAGATC**CCGATTACTAGGCTCCTTAGAC |
| Right thumb nail | *WDR76* | AGAT**CTTGTA**CCGATTACTAGGCTCCTTAGAC |
| Right toe nail | *WDR76* | AGAT**TTAGGC**CCGATTACTAGGCTCCTTAGAC |
| Buccal swab | *WDR76* | AGAT**TAGCTT**CCGATTACTAGGCTCCTTAGAC |
| Duodenum | *WDR76* | AGAT**AGTTCC**CCGATTACTAGGCTCCTTAGAC |
| Colon | *WDR76* | AGAT**ATGTCA**CCGATTACTAGGCTCCTTAGAC |
| Urine | *WDR76* | AGAT**GTAGAG**CCGATTACTAGGCTCCTTAGAC |

BM, bone marrow; uHSCT, umbilical cord hematopoietic stem cell transplantation.

Underlined 6 bases are the barcodes, following 4-based linker sequences.

**Supplemental Table 8.** Primary antibodies used in this study.

| Target | Company | Cat. No |
|---|---|---|
| EFL1 | Pierce | PA5-23877 |
| SBDS | Abcam | ab154222 |
| eIF6 | Santa cruz | sc-366167 |
| FLAG | Sigma | F3165 |
| a-tubulin | Santa cruz | sc-69969 |
| Lamin B | Santa cruz | sc-6216 |
| B-actin | Abcam | ab6276 |
| GAPDH | Abcam | ab8245 |

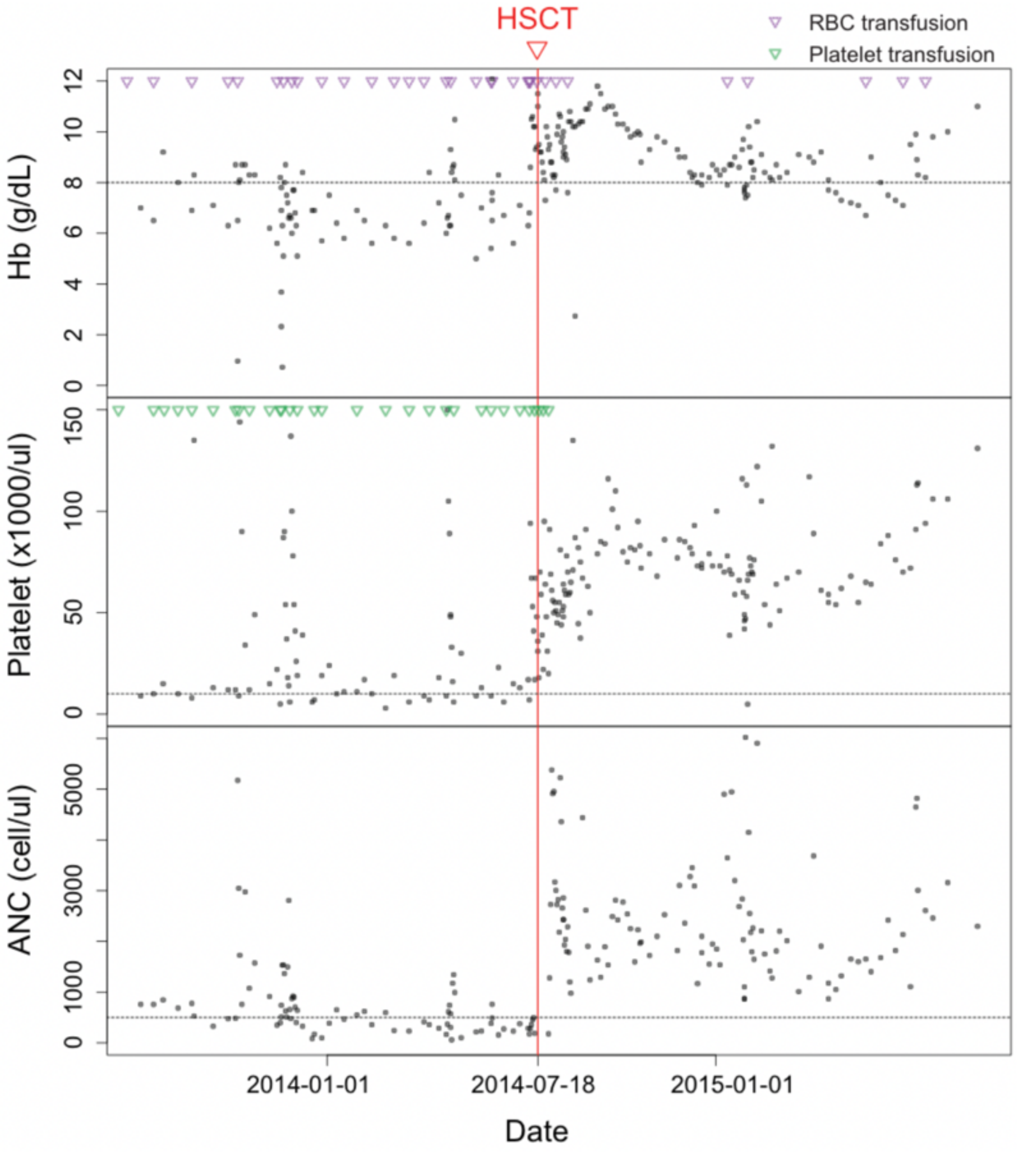


**Supplemental Figure 1.** Hematologic profiles and transfusions before and after hematopoietic stem cell transplantation of I-1. ANC; absolute neutrophil count, Hb; hemoglobin, HSCT; hematopoietic stem cell transplantation, RBC; red blood cell.

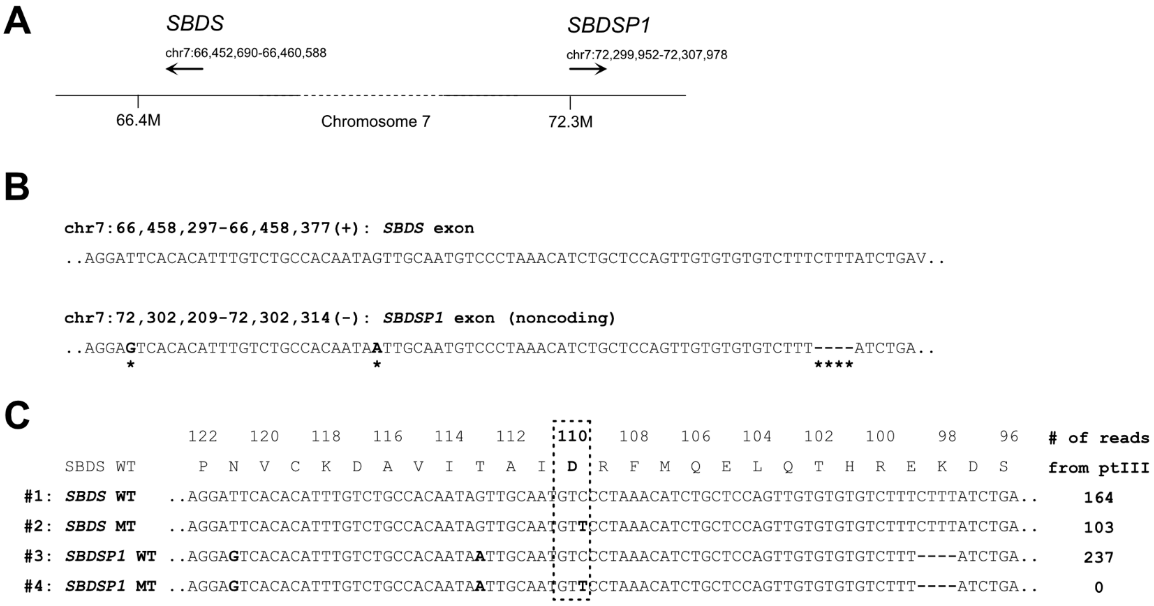


**Supplemental Figure 2.** Validation of *de novo SBDS*$^{p.Asn110Asp}$ variant in III-1 by Illumina sequence reads. (**A**) Locations of *SBDS* and its pseudogene *SBDSP1* on chromosome 7 (hg19). (**B**) Sequence similarity of *SBDS* exon and homologous *SBDSP1* locus encompassing D110 residue. The different bases are indicated by asterisks under the *SBDSP1* sequence. (**C**) Illumina sequence read sequences and number of different read types from patient III-1. Note that number of reads that supports *SBDSP1* variant is zero.

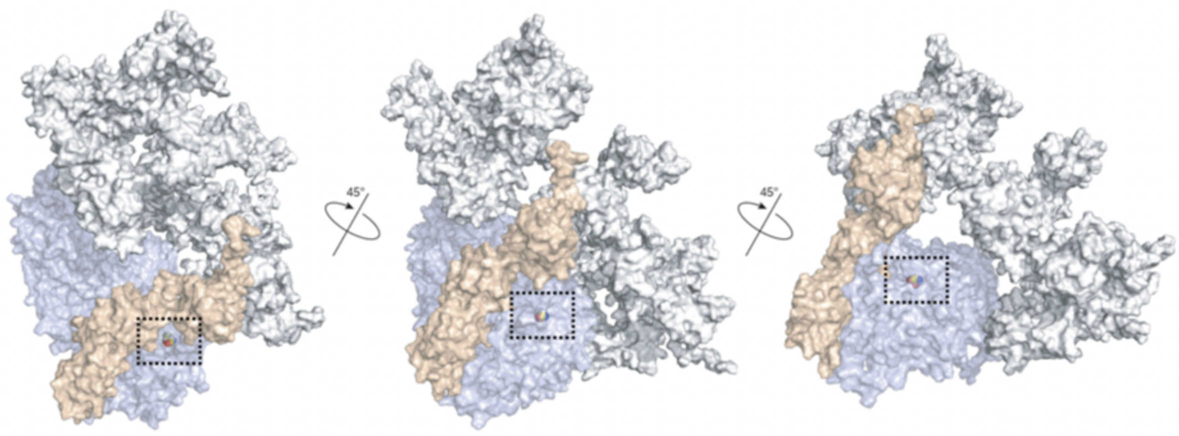


**Supplemental Figure 3.** Structure of EFL1 (violet)-SBDS (orange)-80S (grey) complex, revealing the p.Thr1069 residue being buried inside of EFL1, rather than engaged in SBDS interaction. The p.Thr1069 residue is marked in a space-filling model with carbon atoms in yellow, nitrogen in blue and hydrogen in gray colors.

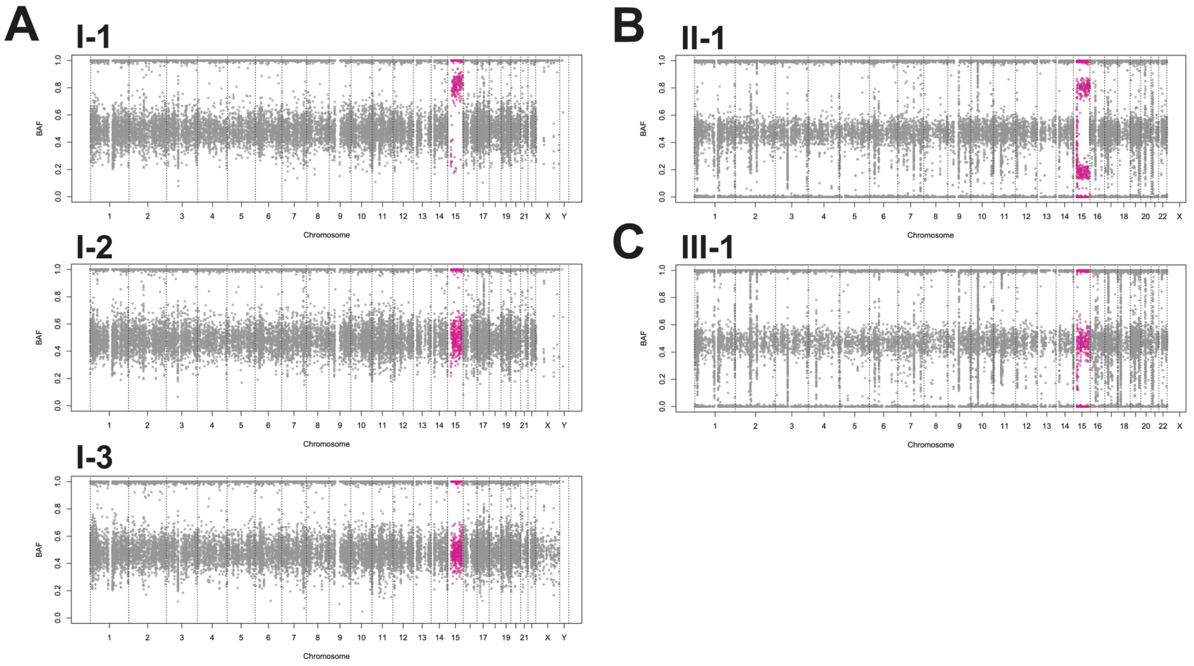


**Supplemental Figure 4.** B-allele frequency (BAF) distributions of variants detected by WES. (**A**) Family I. (**B**) II-1. (**C**) III-1. Chromosome 15 is indicated in red.

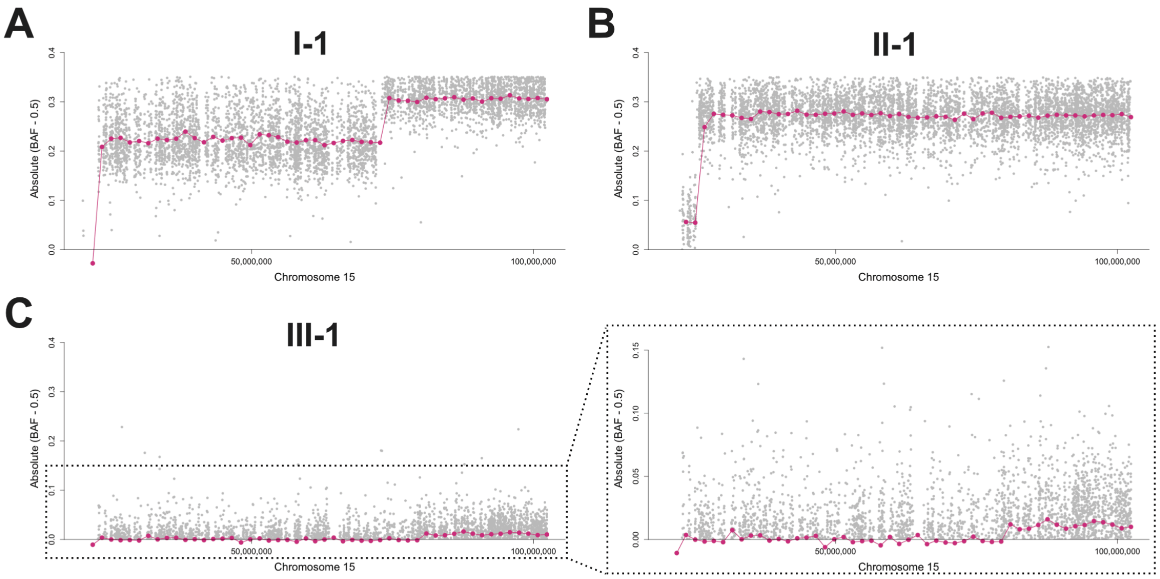


**Supplemental Figure 5.** Degree of loss-of-heterozygosity (LOH) in chromosome 15 depicted by binned average of absolute value of (B allele frequency – 0.5). (**A**) I-1. (**B**) II-1. (**C**) III-1. Right panel shows a zoomed-in view. Gray: each SNP in SNP microarray. Violet: average of absolute value of (B allele frequency – 0.5) binned by 1.6 Mb.

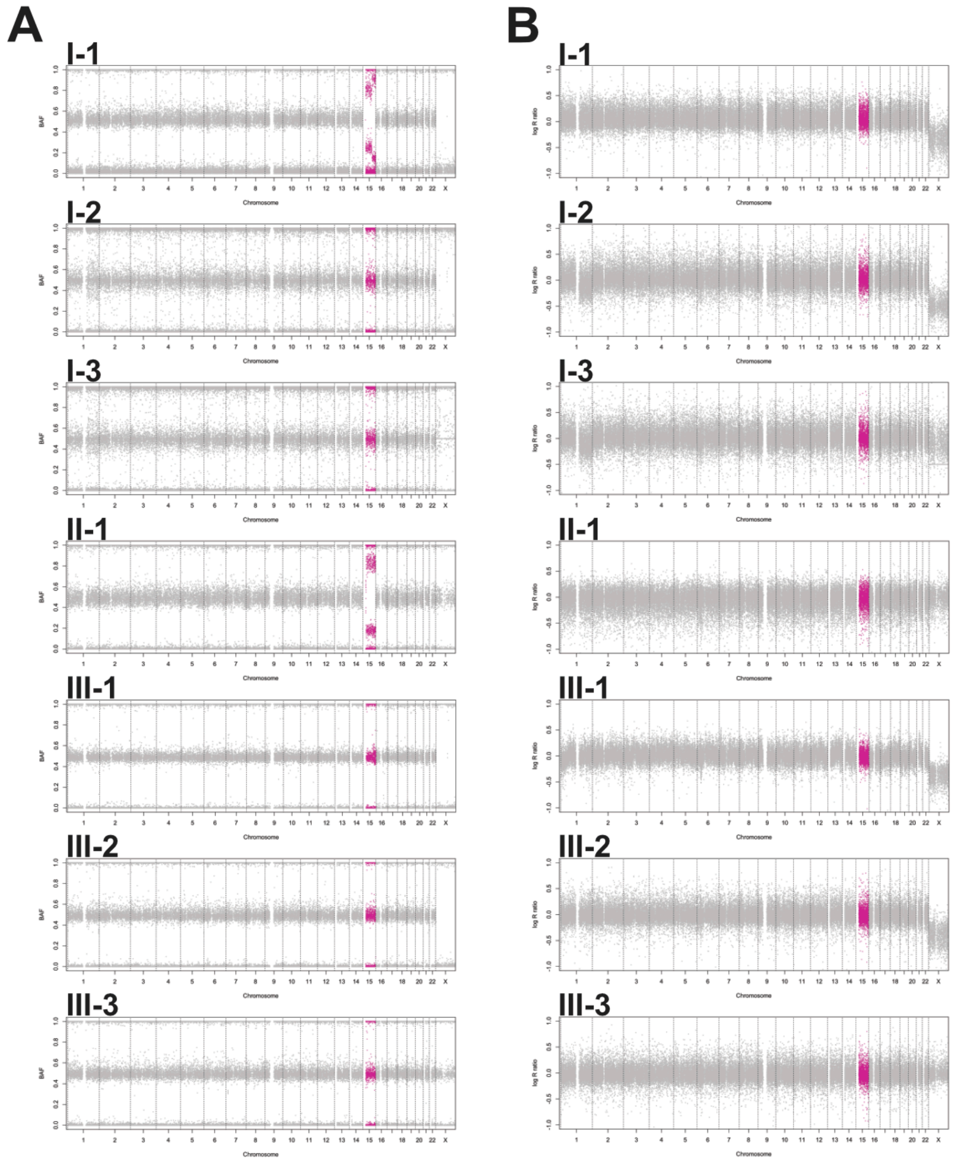


**Supplemental Figure 6.** Genome-wide B-allele frequency and log R ratio by SNP microarray. (**A**) B-allele frequency. (**B**) Log R ratio. Chromosome 15 is indicated in red.

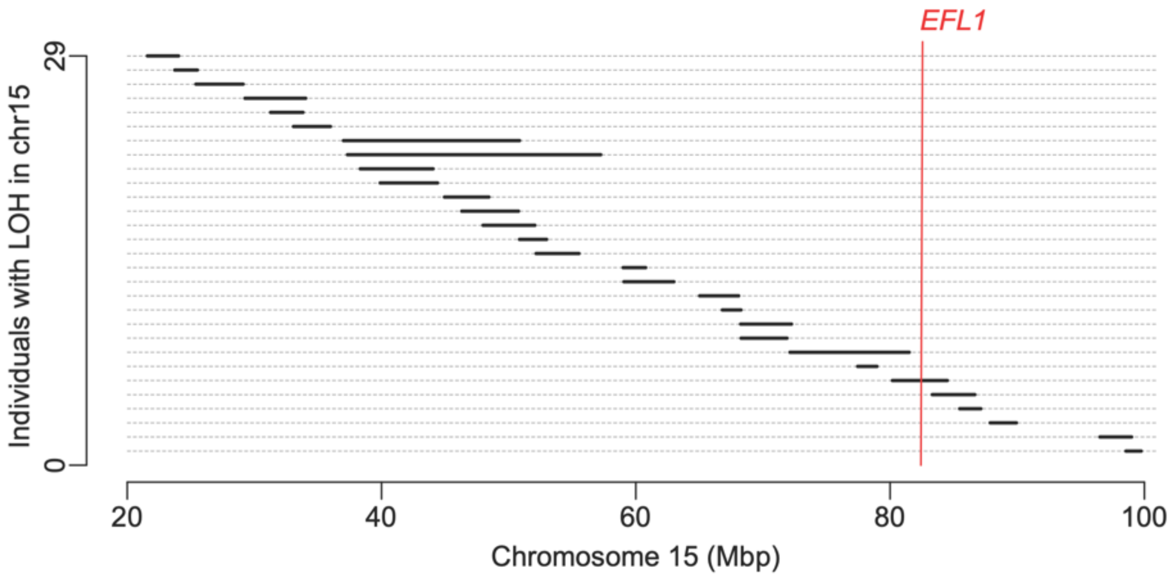


**Supplemental Figure 7.** Chromosome 15 LOH profile of healthy Korean individuals. The SNP array data from 3,667 healthy Korean individuals who participated in the KoGES study (Korean Genome and Epidemiology Study) (Kim et al., 2017) were used to call LOH intervals in chromosome 15 using the PLINK software (http://zzz.bwh.harvard.edu/plink/). 29 individuals carried noticeable LOHs in chromosome 15 and one individual had a LOH surrounding the *EFL1* locus.

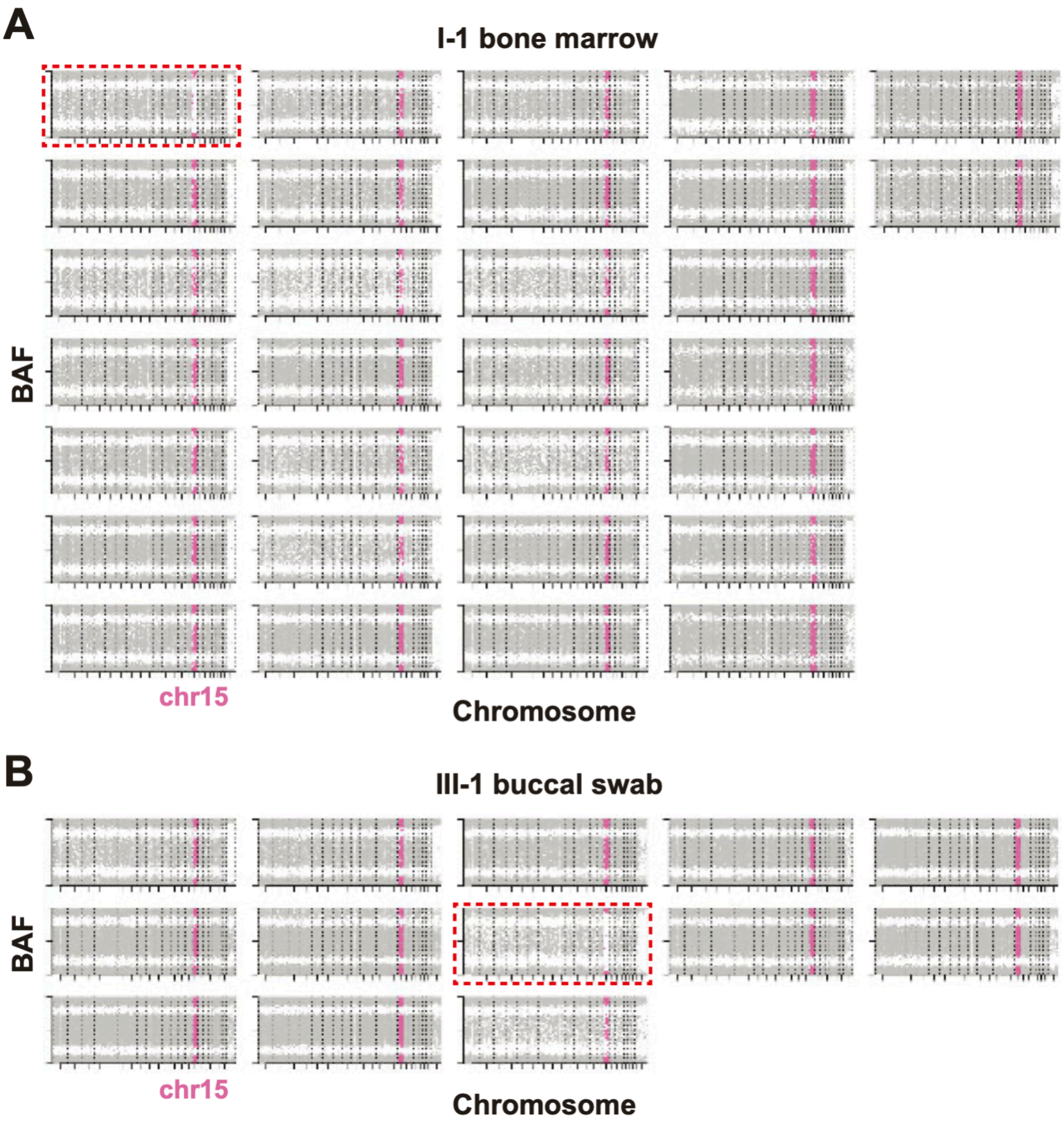


**Supplemental Figure 8.** Whole genome amplified single cell SNP microarray. Each plot displays genome-wide BAF from a single cell. (**A**) SNP array results of 30 whole genome amplified single cells from bone marrow of I-1 among which a cell has complete LOH in chromosome 15 indicating UPD (red rectangle). (**B**) SNP array results of 13 whole genome amplified single cells from buccal swab of III-1 showing that there is a complete LOH in chromosome 15, caused by an UPD event (red rectangle). Chromosome 15 is depicted with pink.

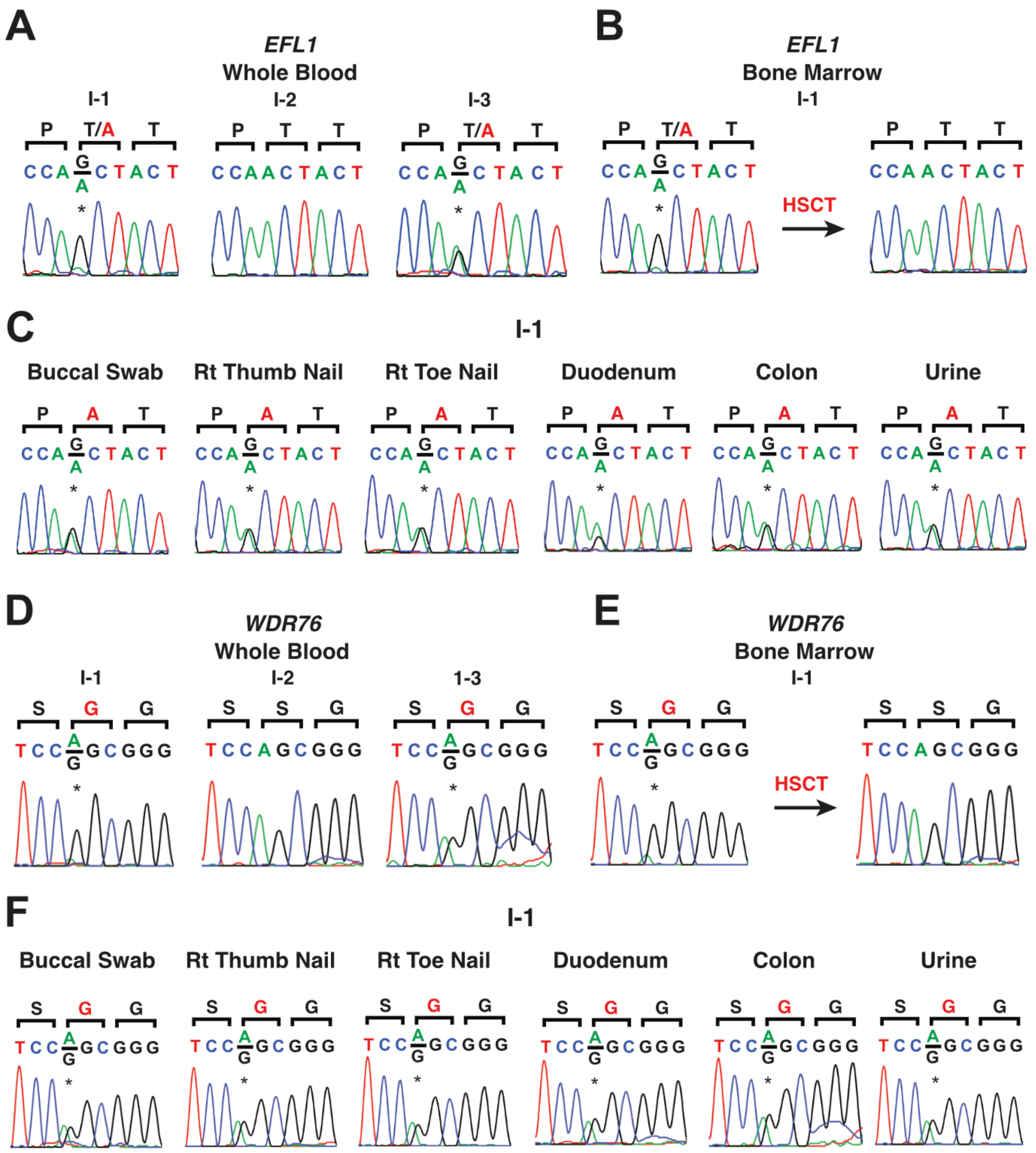


**Supplemental Figure 9.** Sanger sequencing of the *EFL1* (chr15:82,422,872 T>C, NM_024580.5:c.3205A>G) (**A-C**) and *WDR76* variant (chr15:44,158,549 A>G, NM_001167941.1:c.1648A>G) (**D-F**) of family I. the *WDR76* variant was used as a non-functional control. (**A**) Sanger sequencing result of the trio. (**B**) Conversion of the T>C variant to the wild-type allele following allogenic hematopoietic stem cell transplantation (HSCT). (**C**) Sanger sequencing of multiple tissues. (**D**) Sanger sequencing result of the trio. (**E**) Conversion of the T>C variant to the wild-type allele following allogenic hematopoietic stem cell transplantation (HSCT). (**F**) Sanger sequencing of multiple tissues. Rt; right.

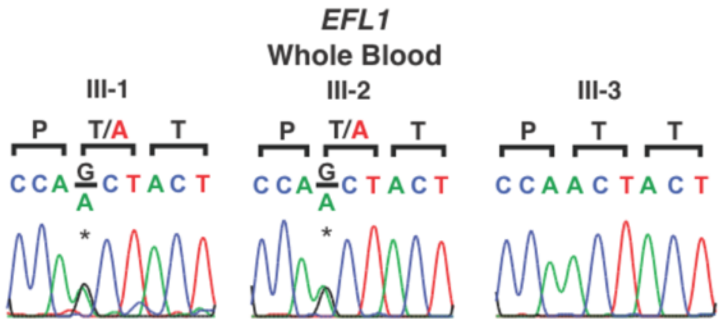


**Supplemental Figure 10.** Sanger sequencing of the *EFL1* variant (chr15:82,422,872 T>C, NM_024580.5:c.3205A>G) of family III.

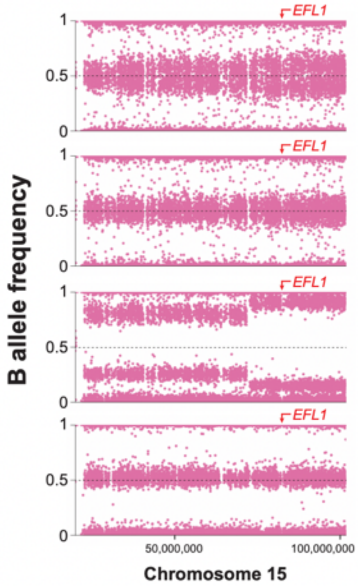


| WBC (x1000 /uL) | Hb (g/dL) | Plt (x1000 /uL) | ANC (cells /uL) | Date of bone marrow exam |
|---|---|---|---|---|
| 5.75 | 5.7 | 3 | 351 | 2011-10-22 |
| 6.08 | 6.2 | 29 | 61 | 2012-03-28 |
| 4.77 | 5.4 | 9 | 382 | 2014-06-05 |
| 3.67 | 8.8 | 72 | 1989 | 2014-10-23 (after HSCT) |

**Supplemental Figure 11.** Change in degrees of mosaicism by time in bone marrow of I-1.

WBC; white blood cells, Hb; hemoglobin, Plt; platelets, ANC; absolute neutrophil count.

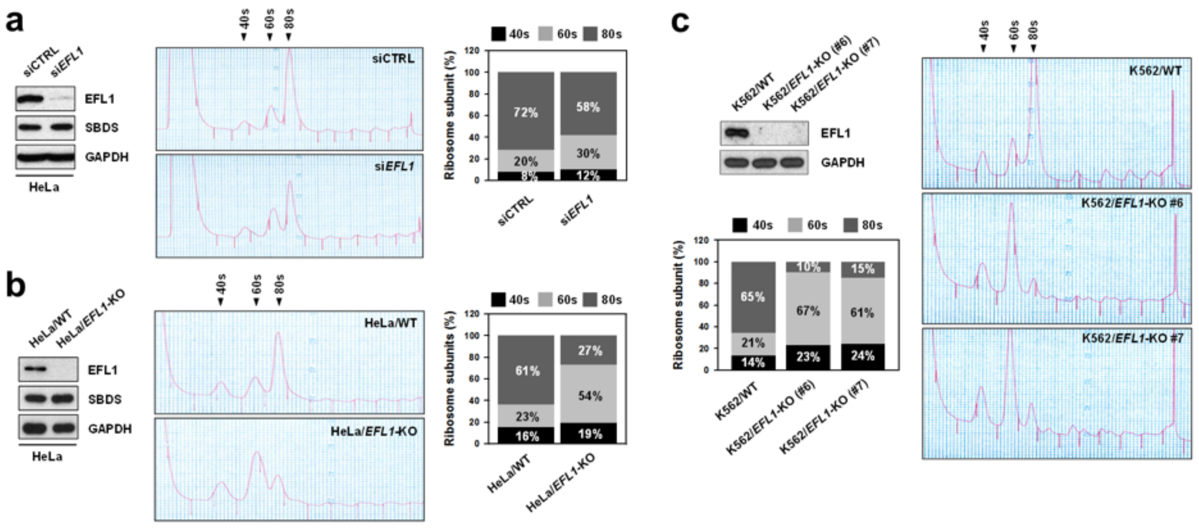


**Supplemental Figure 12.** Polysome profiling when *EFL1* is knocked down by siRNA or knocked-out by CRISPR/Cas9. (**A**) siEFL1 significantly decreased *EFL1* expression and 80S ribosome peak in HeLa cell. (**B-C**) *EFL1* is completely ablated by CRISPR/Cas9 and 80S peak is significantly decreased in HeLa (**B**) and K562 (**C**) cells.

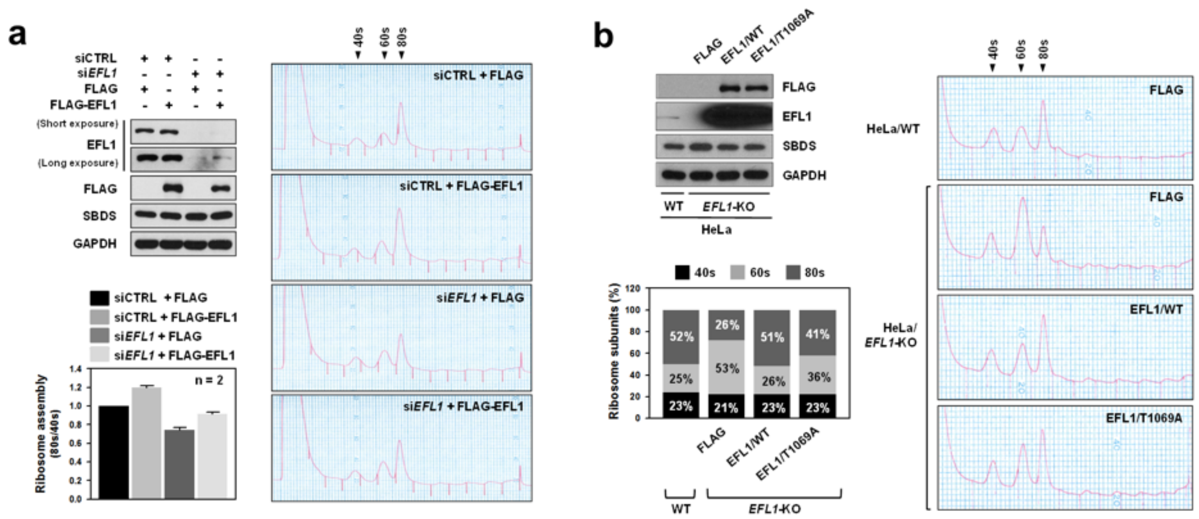


**Supplemental Figure 13.** *EFL1* rescue experiment in HeLa cell. (**A**) 80S ribosome peak is rescued by introduction of wild-type *EFL1* into *EFL1* knocked down HeLa cells. (**B**) Overexpression of wild-type *EFL1* in *EFL1*$^{-/-}$ HeLa cells rescued 80S peak, whereas introduction of *EFL1*$^{p.Thr1069Ala}$ partially did.

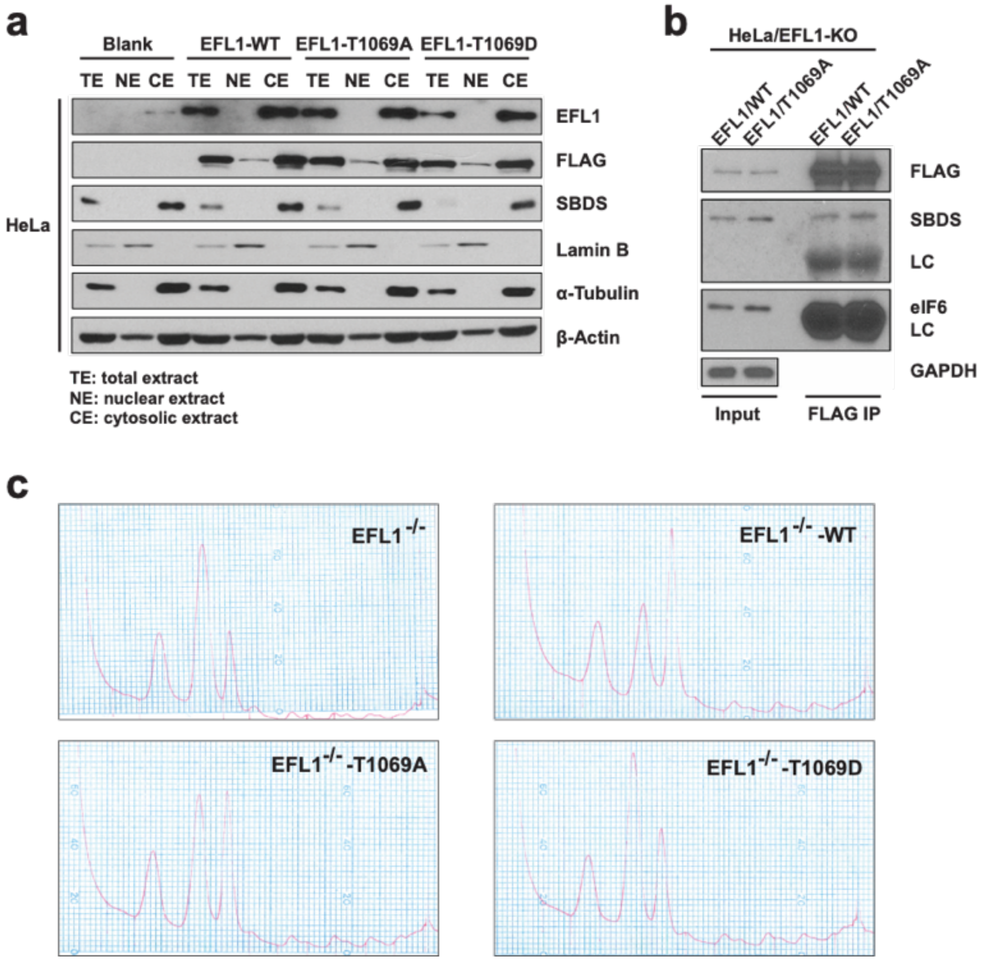


**Supplemental Figure 14.** Seeking for possible mechanism of EFL1 p.Thr1069Ala. (**A**) Subcellular localization of EFL1 is not affected by either p.Thr1069Ala or p.Thr1069Asp overexpressing HeLa cells. (**B**) p.Thr1069Ala mutation does not alter EFL1-SBDS interaction. (**C**) Introduction of EFL1 p.Thr1069Asp into EFL1 KO cell cannot rescue 80S ribosomal peak. p.Thr1069Asp was used as a proxy of phosphorylated Thr1069.

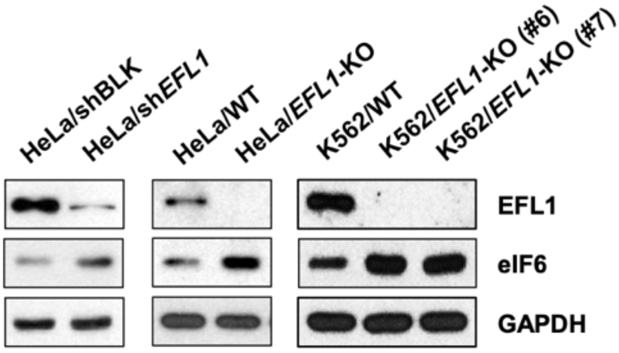


**Supplemental Figure 15**. EFL1 knock-down or knock-out in HeLa and K562 cells increase eIF6 level.

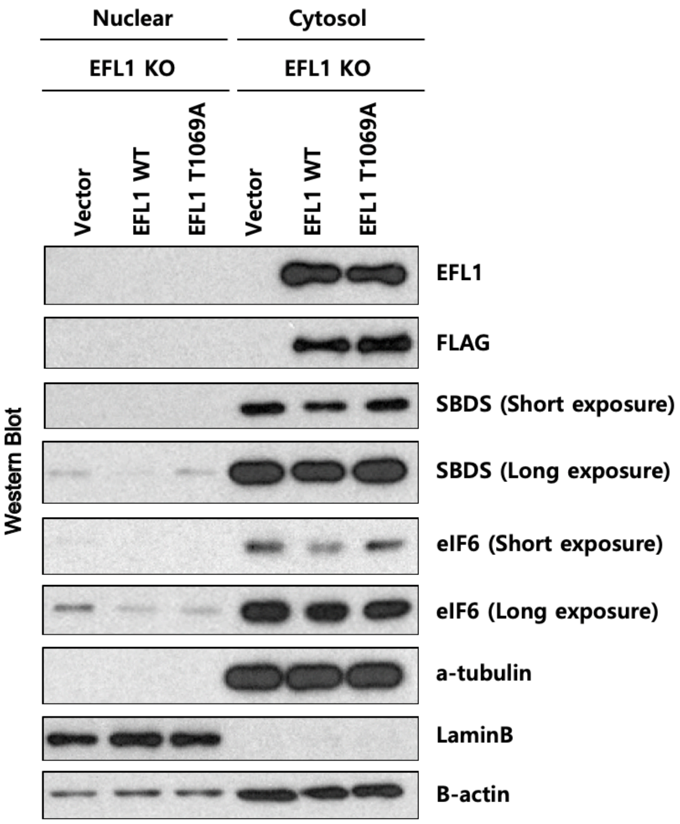


**Supplemental Figure 16.** Nuclear shuttling of eIF6 is impaired by EFL1 knockout or EFL1 p.Thr1069Ala.

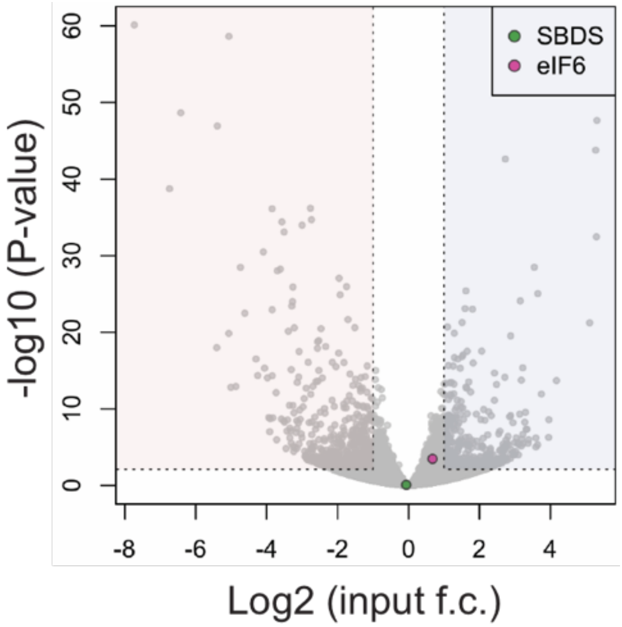


**Supplemental Figure 17.** *SBDS* and *eIF6* expression changes in RNA sequencing of input RNA, demonstrating insignificant changes by the absence of *EFL1*.

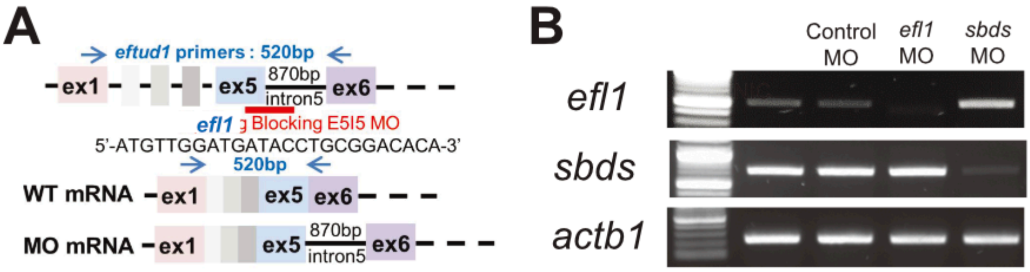


**Supplemental Figure 18.** *efl1* ablation in a zebrafish model. (**A**) Design of MO blocking splicing at exon 5 of *efl1*. (**B**) Knockdown of *efl1* confirmed by reverse transcription PCR. MO; morpholino.

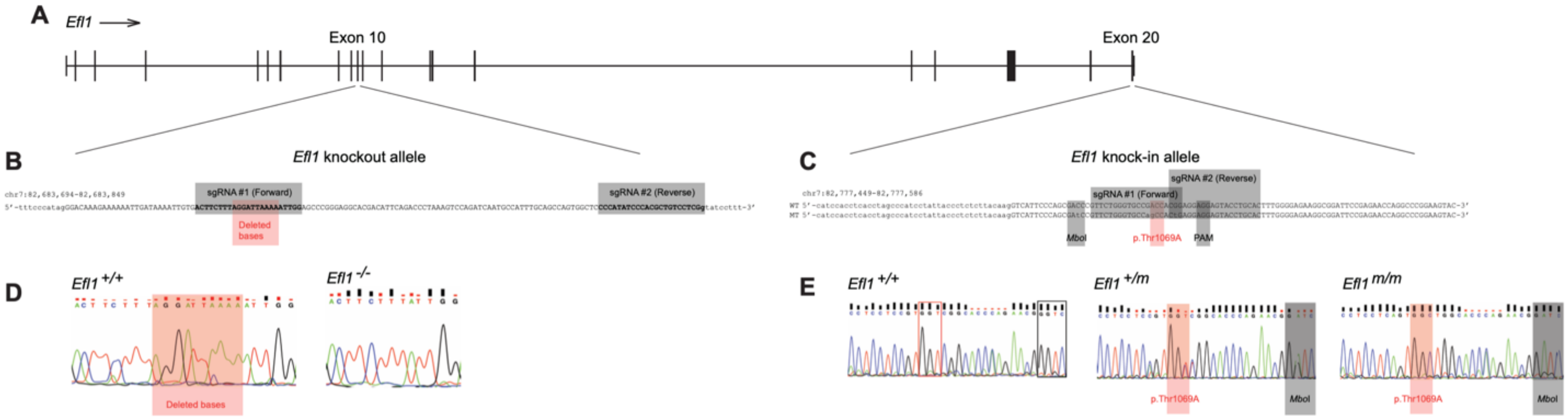


**F** $P = 1.5 \times 10^{-9}$

| Stage | *Efl1* knockout allele +/+ | +/- | -/- | Total |
|---|---|---|---|---|
| P7 | 31 | 54 | 0 | 85 |
| E10.5 | 3 | 6 | 0 | 9 |
| E9.5 | 1 | 8 | 0 | 9 |
| E8.5 | 5 | 8 | 0 | 13 |
| Total | 40 | 76 | 0 | 116 |

**G** $P = 0.76$

| Stage | *Efl1* knock-in allele +/+ | +/m | m/m | Total |
|---|---|---|---|---|
| P30 | 23 | 58 | 24 | 105 |

**H** $P = 0.76$

| *Efl1* knock-in allele | P7 | *Efl1* knockout allele + | - | Total |
|---|---|---|---|---|
| | + | 38 | 49 | 87 |
| | m | 42 | 36 | 78 |
| | Total | 80 | 85 | 165 |

**Supplemental Figure 19.** Construction of *Efl1* mouse models. (**A**) *Efl1* gene structure (mm10). (**B**) Scheme of *Efl1* knockout allele using two sgRNAs. (**C**) Scheme of *Efl1* knock-in allele using two sgRNAs. *Mbo*I locus was introduced for easy genotyping. (**D**) Sanger validation of *Efl1* knockout allele. (**E**) Sanger validation of *Efl1* knockout allele. Note the sequences are in reverse orientation. (**E-G**) Punnett squares of *Efl1* knockout x knckout (**E**), *Efl1* knock-in x knock-in (**F**) and *Efl* CH strain (**G**). *P*-values are Fisher's exact test results of expected and observed offspring numbers of each genotype.

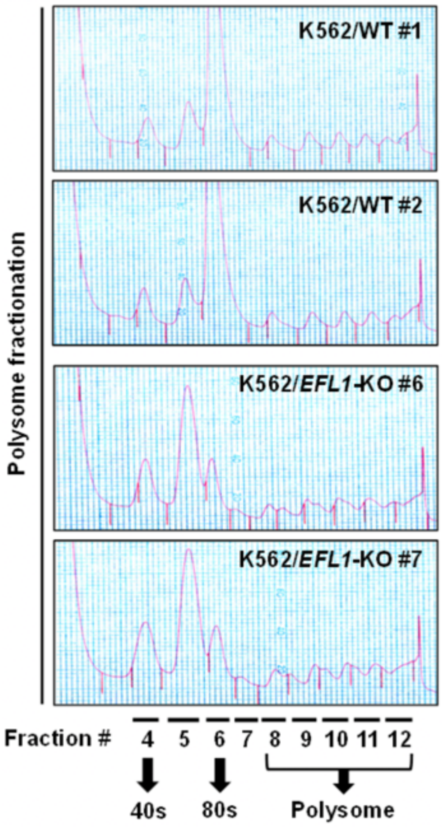


**Supplemental Figure 20.** Polysome fractionation utilized for the RNA sequencing experiment.

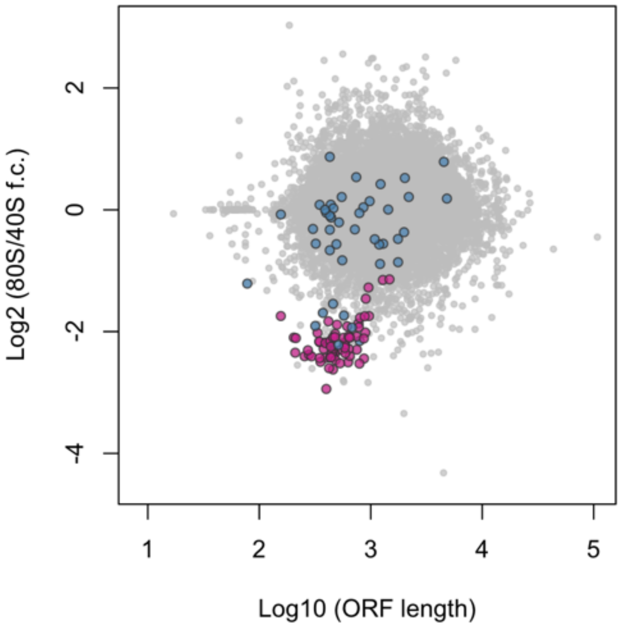


**Supplemental Figure 21.** Fold change of 80S enriched genes normalized by 40S plotted by transcript ORF length. Red; significantly downregulated RP genes in Fig. 2c, blue; other RP genes.

<table>
<tr><th>Gene name</th><th>Start</th><th>p-value</th><th>Sites</th></tr>
<tr><td>ENSG00000177600.4|RPLP2</td><td>318</td><td>3.22e-8</td><td>GCGUCGGCGC CUUCCUUUUCCUCCC UGUCGCCACC</td></tr>
<tr><td>ENSG00000143947.8|RPS27A</td><td>283</td><td>2.89e-7</td><td>GCGCGGCGUU CUUCCUUUUCGAUCC GCCAUCUGCG</td></tr>
<tr><td>ENSG00000156482.6|RPL30</td><td>458</td><td>3.77e-7</td><td>CGCCGUCAUU CUUUUUUUUCCUUCU UUGAGACAGA</td></tr>
<tr><td>ENSG00000174748.14|RPL15</td><td>600</td><td>5.97e-7</td><td>GAGUACAGCU CUUUCCUUUCCGUCU GGCGGCAGCC</td></tr>
<tr><td>ENSG00000142676.8|RPL11</td><td>24</td><td>9.52e-7</td><td>GGAAGCUCCG CUUUCUCUUCCUGCU CUCCAUC</td></tr>
<tr><td>ENSG00000118181.6|RPS25</td><td>344</td><td>1.19e-6</td><td>CUGAGCAGCG CUUCCUUUUUGUCCG ACAUCUUGAC</td></tr>
<tr><td>ENSG00000114391.8|RPL24</td><td>65</td><td>2.71e-6</td><td>AUGAUUCUCU CUUUCUUUUCGCCAU CUUUUGUCUU</td></tr>
<tr><td>ENSG00000122026.6|RPL21</td><td>246</td><td>3.93e-6</td><td>GGCCCCGCCU CUUUCCUUUCGGCCG GAACCGCCAU</td></tr>
<tr><td>ENSG00000197756.5|RPL37A</td><td>652</td><td>5.60e-6</td><td>CACCGCGUCU CUUCCUUUCUGGGCU CGGACCUAGG</td></tr>
<tr><td>ENSG00000110700.2|RPS13</td><td>117</td><td>7.92e-6</td><td>UCCACUCUCG CUCUCCUUUCGUUGC CUGAUCGCCG</td></tr>
<tr><td>ENSG00000161970.8|RPL26</td><td>3</td><td>9.33e-6</td><td>CU CUUCCCUUUUGCGGC CAUCACCGAA</td></tr>
<tr><td>ENSG00000198918.7|RPL39</td><td>21</td><td>1.09e-5</td><td>CUUCCCUCCU CUUCCUUUCUCCGCC AUCGUGGUGU</td></tr>
<tr><td>ENSG00000122406.8|RPL5</td><td>51</td><td>1.49e-5</td><td>AUGCAUGGAG GUUCCCUUUUCUUGC CCGUAUGCCA</td></tr>
<tr><td>ENSG00000138326.14|RPS24</td><td>102</td><td>1.74e-5</td><td>CGAAUCGUGG UUCUCUUUUCCUCCU UGGCUGUCUG</td></tr>
<tr><td>ENSG00000147403.12|RPL10</td><td>538</td><td>1.74e-5</td><td>AGGAGCGCCU CUUUCCCUUCGGUGU GCCACUGAAG</td></tr>
<tr><td>ENSG00000168028.9|RPSA</td><td>28</td><td>1.74e-5</td><td>UUCCUGCCGC CUGUCUUUUCCGUGC UACCUGCAGA</td></tr>
<tr><td>ENSG00000166441.8|RPL27A</td><td>27</td><td>3.07e-5</td><td>UCGUUUCUUG CCUCCUCUUCCUUGC CGGCGGAGAC</td></tr>
<tr><td>ENSG00000161016.11|RPL8</td><td>194</td><td>3.07e-5</td><td>UGACGCCGUG UUUCCUCUUUCGGCC GCGCUGGUGA</td></tr>
<tr><td>ENSG00000145592.9|RPL37</td><td>111</td><td>3.52e-5</td><td>GCGGAAGUGC CUUCUCUUCCGGUCU UUCUGGUCUC</td></tr>
<tr><td>ENSG00000142937.7|RPS8</td><td>135</td><td>4.02e-5</td><td>AUCUUUGCGG UUUCUCUUUCCAGCC AGCGCCGAGC</td></tr>
<tr><td>ENSG00000140988.11|RPS2</td><td>161</td><td>4.02e-5</td><td>ACACUGACUA GUUCCUUCUUGUCGC UUUUCCCAGC</td></tr>
<tr><td>ENSG00000164587.7|RPS14</td><td>18</td><td>4.58e-5</td><td>CCCUCCCACU CUCUCUUUCCGGUGU GGAGUCUGGA</td></tr>
<tr><td>ENSG00000071082.6|RPL31</td><td>547</td><td>7.51e-5</td><td>GCAACUGCGG CUUUCCUUCUCCCAC AAUCCUUCGC</td></tr>
<tr><td>ENSG00000198242.9|RPL23A</td><td>76</td><td>7.51e-5</td><td>CAAAUGUGUU CUUUCCUCUUGGCAG CGUUCCUGCC</td></tr>
<tr><td>ENSG00000125691.8|RPL23</td><td>268</td><td>7.51e-5</td><td>GCUGAACAGA CUUCCCGCUGCUGGU GUGGGUGAC</td></tr>
<tr><td>ENSG00000105640.8|RPL18A</td><td>520</td><td>9.49e-5</td><td>CUCACUCCUG UUUCCUUUCAGCUAC GAGAGUACAA</td></tr>
<tr><td>ENSG00000134419.11|RPS15A</td><td>240</td><td>9.49e-5</td><td>AGCGUCUGGA UCUCCCUUUAGUGCU GAGGCAGCGC</td></tr>
<tr><td>ENSG00000204628.7|GNB2L1</td><td>131</td><td>1.06e-4</td><td>UCGGCCUACU UUUUCUGCCCCUGGU UCUCCCCGUC</td></tr>
<tr><td>ENSG00000148303.12|RPL7A</td><td>80</td><td>1.06e-4</td><td>CCUCGGAGGG CCUCCUGCUCCUCCU GGCGCUAGGA</td></tr>
<tr><td>ENSG00000131469.8|RPL27</td><td>50</td><td>1.06e-4</td><td>CACUGUUCUG UUUCUCUUUGGUCAU UUCUCAUAGC</td></tr>
<tr><td>ENSG00000089009.11|RPL6</td><td>123</td><td>1.19e-4</td><td>CUUCCAGACG CUUCAUUUUUGUUGU UUGGGUUUGC</td></tr>
<tr><td>ENSG00000105372.2|RPS19</td><td>172</td><td>1.33e-4</td><td>CCCCUUGGCC CUCCCCUUUACCUCU CCACCCCUCA</td></tr>
<tr><td>ENSG00000142534.2|RPS11</td><td>1</td><td>1.64e-4</td><td>CUUUUUUUCAGGCGG CCGGGAAGAU</td></tr>
<tr><td>ENSG00000142541.12|RPL13A</td><td>51</td><td>1.64e-4</td><td>GACAAAACCU CCUCCUUUUCCAAGC GGCUGCCGAA</td></tr>
<tr><td>ENSG00000137818.7|RPLP1</td><td>38</td><td>1.82e-4</td><td>GCGAGAGCCC CUUUCCUCAGCUGCC GCCAAGGUGC</td></tr>
<tr><td>ENSG00000108298.5|RPL19</td><td>151</td><td>1.82e-4</td><td>GGCCGGAGCA CUUUCGUCCCGGGCC UCCGGAGUGA</td></tr>
<tr><td>ENSG00000186468.8|RPS23</td><td>222</td><td>2.02e-4</td><td>UGCGGUGCUU CUCUCUUUCGCUCAG GCCCGUGGCG</td></tr>
<tr><td>ENSG00000182899.10|RPL35A</td><td>203</td><td>2.23e-4</td><td>GGGGCCUCGU CCUUCUCUUACCGCC AUCUUGGCUC</td></tr>
<tr><td>ENSG00000144713.8|RPL32</td><td>251</td><td>2.23e-4</td><td>GAGCUGCGGG CUGCUUGUUUGUUCC UUGGCGGUGG</td></tr>
<tr><td>ENSG00000171490.8|RSL1D1</td><td>93</td><td>2.23e-4</td><td>CUCGGCCUCG CUGUCUUCUGCAGCC GCUACUGGAA</td></tr>
<tr><td>ENSG00000198034.6|RPS4X</td><td>80</td><td>2.46e-4</td><td>GAAGACGGAG GUCCUCUUUCCUUGC CUAACGCAGC</td></tr>
<tr><td>ENSG00000171863.8|RPS7</td><td>49</td><td>3.27e-4</td><td>CGGUUUCCGC CCUCCUCCUCGCGCU GUUUCCGCCU</td></tr>
<tr><td>ENSG00000116251.5|RPL22</td><td>548</td><td>3.27e-4</td><td>CAUCAGCACA GUCUCUUCUGGUGAU UGGACUUUUG</td></tr>
<tr><td>ENSG00000147604.9|RPL7</td><td>43</td><td>3.58e-4</td><td>CCUCCUCGCA CCUCCCUUCGCAGCU CACGCCCCCU</td></tr>
<tr><td>ENSG00000008988.5|RPS20</td><td>50</td><td>3.92e-4</td><td>UGGUCCGCAC GCUCCUGCUCCUGAC UCACCGCUGU</td></tr>
<tr><td>ENSG00000089157.11|RPLP0</td><td>296</td><td>4.29e-4</td><td>CUUACUUGGA CUUCCUUUCCUGUAG GGAAUCUCAC</td></tr>
<tr><td>ENSG00000112306.7|RPS12</td><td>127</td><td>4.67e-4</td><td>UGCGGAUGAG GCCUCUUUCCCUGCC GCCGCCGAGU</td></tr>
<tr><td>ENSG00000188846.9|RPL14</td><td>31</td><td>5.09e-4</td><td>CAUGGUGAGU CUUACUGUUGCGGGC UCCGGGGCCG</td></tr>
<tr><td>ENSG00000137154.8|RPS6</td><td>469</td><td>6.01e-4</td><td>GCAUCUUAUA UUUCCUGCUUGUGUG GAGGCAACAU</td></tr>
<tr><td>ENSG00000177954.7|RPS27</td><td>1</td><td>6.01e-4</td><td>GCUCCUUUCCGGCGG UGACGACCUA</td></tr>
<tr><td>ENSG00000174444.10|RPL4</td><td>208</td><td>6.52e-4</td><td>CACUUGCCAC CUUCCAUCUCGUUUG AUGACGUACA</td></tr>
<tr><td>ENSG00000167526.9|RPL13</td><td>2</td><td>6.52e-4</td><td>C CUUUCCGCUCGGCUG UUUUCCUGCG</td></tr>
<tr><td>ENSG00000231500.2|RPS18</td><td>63</td><td>7.06e-4</td><td>CUGUGUCUGA UUUCUUCCCCCGUAC UUUUUCAACU</td></tr>
<tr><td>ENSG00000213741.4|RPS29</td><td>1</td><td>7.06e-4</td><td>CUUUUACCUCGUUGC ACUGCUGAGA</td></tr>
<tr><td>ENSG00000170889.9|RPS9</td><td>117</td><td>7.64e-4</td><td>GCUUGCGCGC CUCUUUCUCAGUGAC CGGGUGGUUU</td></tr>
<tr><td>ENSG00000105193.4|RPS16</td><td>3</td><td>8.25e-4</td><td>CU UUUCCGGUUGCGGCG CCGCGCGGUG</td></tr>
<tr><td>ENSG00000149273.10|RPS3</td><td>26</td><td>8.89e-4</td><td>ACUUGGGGAU GUUCUCUUUGCCCAG GUGGCCUACU</td></tr>
<tr><td>ENSG00000163682.11|RPL9</td><td>278</td><td>1.03e-3</td><td>CGCUUUGGGG CUGCUCCUUCCAGGC UGGGACACAC</td></tr>
<tr><td>ENSG00000063177.8|RPL18</td><td>474</td><td>1.11e-3</td><td>CAGUGUCUGG CUGUUCGUUGGGUAC ACAGUAAUCU</td></tr>
<tr><td>ENSG00000198755.6|RPL10A</td><td>1</td><td>1.27e-3</td><td>UUUCCGGUUAGCGCG GCGUGAGAAG</td></tr>
<tr><td>ENSG00000221983.3|UBA52</td><td>558</td><td>1.46e-3</td><td>CUCGAGCGAU UCUUCUGCCUCAGCC UCCCGAGUAG</td></tr>
<tr><td>ENSG00000172809.8|RPL38</td><td>205</td><td>1.77e-3</td><td>CCGAGGACUG UUUCCCGCAGGUCCU GGUCCGCGCC</td></tr>
<tr><td>ENSG00000197958.8|RPL12</td><td>8</td><td>1.88e-3</td><td>CUCUCGG CUUUCGGCUCGGAGG AGGCCAAGGU</td></tr>
<tr><td>ENSG00000109475.12|RPL34</td><td>85</td><td>2.39e-3</td><td>GUAGGGCGGU GUUUCUCCAACUGGG CUCGUGGCAC</td></tr>
<tr><td>ENSG00000233927.4|RPS28</td><td>1</td><td>2.98e-3</td><td>CUCUCCGCCAGACCG CCGCCGCGCC</td></tr>
<tr><td>ENSG00000162244.6|RPL29</td><td>98</td><td>4.64e-3</td><td>GACCGACCGU GUGUCUGUCGGGAGC AAAAGCUGCG</td></tr>
</table>

**Supplemental Figure 22.** Enriched motif in 5' UTR of RP genes that were decreased in the 80S-bound compared to the 40S-bound fraction in the mutant cells. Motif sequences and their location in 5' UTR of RP genes.
